# Werner syndrome RECQ helicase participates in and directs maintenance of the protein complexes of constitutive heterochromatin in proliferating human cells

**DOI:** 10.1101/2024.01.29.577850

**Authors:** Pavlo Lazarchuk, Matthew Manh Nguyen, Crina M. Curca, Maria N. Pavlova, Junko Oshima, Julia M. Sidorova

## Abstract

The WRN RECQ helicase is responsible for the Werner syndrome of premature aging and cancer predisposition. Substantial progress has been made in delineating WRN functions in multiple aspects of DNA metabolism, including DNA replication, repair, transcription, and telomere maintenance. Nevertheless, a complete mechanistic understanding of how loss of WRN accelerates aging in humans has not been achieved yet. Here we show that WRN is involved in the maintenance of constitutive heterochromatin, CH, in proliferating, immortalized human fibroblasts. WRN is found within a complex with histone deacetylase 2, HDAC2, and WRN/HDAC2 association is mediated by heterochromatin protein alpha, HP1α. WRN deficiency derepresses SATII pericentromeric satellite repeats and reduces a subset of protein-protein interactions that participate in the organization of CH in the nucleus. In particular, WRN deficiency reduces the complexes involving Lamin B1 and Lamin B receptor, LBR. Both mRNA level and subcellular distribution of LBR are affected by WRN deficiency, and the latter phenotype does not require WRN catalytic activities. At the mRNA level, WRN supports complete maturation of the LBR mRNA. All signs of heterochromatin disruption seen in WRN-deficient proliferating fibroblasts are also observed in WRN-proficient fibroblasts undergoing replicative or oncogene-induced senescence, and WRN complexes with HP1α and HDAC2 are also markedly downregulated in these senescing cells. The data suggest that WRN loss affects heterochromatin independently of the senescence program but can mimic aspects of it and thus sensitize cells to triggers of senescence.

## Introduction

The RECQ helicase/exonuclease WRN is responsible for the Werner syndrome of premature aging, WS (Oshima et al., 2017b). WRN is one of the five RECQ helicases in human cells and, as other RECQs, WRN is involved in most aspects of DNA metabolism (Sidorova and Monnat Jr, 2015). In replication, WRN maintains optimal replication fork progression through difficult-to-replicate regions of the genome (Drosopoulos et al., 2015; Murfuni et al., 2012; van Wietmarschen et al., 2020), which at least in some case is due to its ability to unwind unusual DNA structures such as G-quadruplexes, and unwind and degrade RNA:DNA hybrids (Suzuki et al., 1997; Suzuki et al., 1999). Indeed, WRN-deficient and WRN helicase-dead (K577M)-expressing cells have been shown to upregulate nuclear RNA:DNA hybrids (Marabitti et al., 2019) *in vivo*. Nevertheless, WS stands out among other RECQ deficiency syndromes by its preponderance of pathologies found in normal aging yet appearing on an accelerated schedule (Lautrup et al., 2019; Oshima et al., 2017a; Poot, 2017), urging an ongoing search for the unique biochemical functions and/or cellular roles of WRN.

In this light, WRN is the only RECQ helicase thus far implicated in heterochromatin maintenance (Lee et al., 2020; Zhang et al., 2015). Specifically, accelerated senescence of WRN-deficient mesenchymal stem cells in culture was accompanied by symptoms of loss of constitutive heterochromatin, CH, such as reduction of heterochromatic histone mark H3K9me3 and elevated expression of satellite repeat sequences that comprise pericentromeric constitutive heterochromatin, or PCH (Zhang et al., 2015). The study also found that WRN physically interacted with the histone methylase SUV39H1 and heterochromatin protein HP1α (CBX5), and was detected on PCH by ChIP. In line with localization of PCH to the nucleoli or nuclear lamina (Bizhanova and Kaufman, 2021; Brändle et al., 2022; Németh et al., 2010; Padeken and Heun, 2014), WRN is found in the nucleoli (Marciniak et al., 1998) and co-immunoprecipitates with Lamin B1 in a DNA-dependent manner (Lachapelle et al., 2011). Importantly, relaxation of CH and PCH is likely a common feature of aging and senesced cells (Rocha et al., 2021), and it has been argued that WRN deficiency is a case that demonstrates a cause-effect relationship between loss of CH and aging (Zhang et al., 2015). However, many mechanistic questions remain. For example it is unclear if WRN-mediated effect on heterochromatin in MSCs is independent from the replicative senescence program, which limits their replicative lifespan to 20-40 population doublings. Also, despite the findings of physical interaction of WRN with heterochromatin constituents, it is unclear what activity or role WRN contributes to the assembly and functioning of heterochromatin and whether it is distinct and independent of WRN’s other roles. Lastly, if WRN is involved critically and directly in heterochromatin maintenance, it is of interest to determine if WRN or its interactions are targeted by the senescence programs when they operate in WRN-proficient cells.

The deacetylases HDAC1 and HDAC2 are involved in the establishment of repressive chromatin genome-wide, including the PCH (Craig et al., 2003; Helbling Chadwick et al., 2009; Sims and Wade, 2011). HDAC2 was found in complex with SUV39H1 (Vaute et al., 2002) and Lamin A/C (Mattioli et al., 2018), and an early study detected HDAC2 but not HDAC1 in a multiprotein complex specific to senescent cells (Wagner et al., 2001). We originally identified a functional interaction between WRN and HDAC1 at stalled replication forks and found that WRN, HDAC1, and HDAC2, though detectable on replicating DNA, did not move with the replication fork but remained associated with maturing chromatin for hours after replication (Kehrli et al., 2016). In addition, a fraction of WRN co-immunoprecipitated with HDAC2 and HDAC1. WRN presence in the immunoprecipitates by either HDAC1 or HDAC2 antibody was dependent primarily on HDAC2 presence in the lysates. We hypothesized that WRN’s co-immunoprecipitation with HDAC1 is due either to HDAC1 antibody’s cross-reactivity to HDAC2 (due to the proteins’ high level of homology), or to the fact that HDAC1 and HDAC2 can form a complex.

Here, we examined the role of WRN in maintenance of CH in proliferating cells, placing its association with HDAC2 in the context of protein assemblies involving HP1α, Lamin B1, and Lamin B receptor, the assemblies that undergo dramatic downregulation in senescing cells.

## Materials and Methods

### Cells and culture

The SV40-transformed human fibroblast GM639 (GM00639, Cellosaurus ID CVCL_7299) and its derivatives have been used by us previously (Lazarchuk et al., 2020; Shukla et al., 2022; Sidorova et al., 2008) and was originally obtained from the NIGMS Human Genetic Mutant Cell Repository. Early and late passage normal human dermal fibroblasts (NHDFs) were generated in-house and are described in (Kehrli and Sidorova, 2014). GM639 and NHDFs were grown in high glucose Dulbecco’s Modified Minimal Essential Medium (DMEM) with L-glutamine, 10% fetal bovine serum, FBS, (Hyclone) and antibiotics. WI38hTERT fibroblast line was a gift of Dr. Carl Mann and contains an inducible GFP-RAF1-ER transgene (Jeanblanc et al., 2012). WI-38hTERT was grown in Minimal Essential Medium (MEM) with L-glutamine and sodium pyruvate (Gibco) supplemented with 10% FBS, Nonessential Amino Acids, and antibiotics. All cell lines were kept in a humidified 5% CO2, 37°C incubator. Mycoplasma testing was performed using the UW/FHCC Cancer Consortium Shared Resource Specimen processing service https://sharedresources.fredhutch.org/services/mycoplasma-testing.

### Drugs and other reagents

Stock of 5-chlorodeoxyuridine (CldU, Sigma-Aldrich) was at 10mM in PBS, and 5-ethynyldeoxyuridine (EdU, Sigma-Aldrich or Click Chemistry Tools) was at 10mM in DMSO. CldU was used at a concentration of 100μM and EdU was used at 10 or 20μM. Doxycycline stock solution was at 100μg/ml M in PBS and it was used at 100ng/ml final concentration. 4-hydroxytamoxifen, 4-HT, stock (Sigma-Aldrich) was 40μM in ethanol and was used at a final concentration of 20nM. All stocks were stored at −20°C.

### Constructs and CRISPR-Cas9-mediated gene knockout

gRNAs were designed against the *WRN* gene sequences GGTAATATTTAACCTCCGT and GTCTATCCGCTGTAGCAAT. The two gRNAs were cloned into the pLenti-L3US2-RFPv3 vector, and the procedures were as described before (Kehrli et al., 2016; Lazarchuk et al., 2020). Briefly, GM639 cells were transduced with a lentiviral vector pLenti-Cas9-2TA-emGFP, and flow-sorted for positives. These cells, GM639-Cas9, were further transduced with dgRNA-expressing pLenti-L3US2-RFPv3. Transduced mass cultures were sorted for RFP-positive cells, and after expansion the loss of expression of WRN was verified by Western blotting. Individual clones were subsequently derived from this cell culture, Western blot verified, and the regions surrounding gRNA sites were PCR-amplified and sequenced.

Virus generation and cell transduction were as described (Sidorova et al., 2008).

WRN expression constructs pLX209-neo-WRN, pLX209-neo-WRN E84A, pLX209-neo-WRN K577M were a gift from Francisca Vazquez (respectively, Addgene plasmids # 125788, http://n2t.net/addgene:125788; RRID:Addgene_125788; #125789, http://n2t.net/addgene:125789; RRID:Addgene_125789 and # 125790; http://n2t.net/addgene:125790; RRID:Addgene_125790) (Chan et al., 2019)).

pLKO.1-Puro-shWRN2 TET ON and pLKO.1-Puro-shNS TET ON for doxycycline-inducible shRNA expression were a gift of Dr. Weiliang Tang. shWRN2 is identical to shWRN2-4 described in (Sidorova et al., 2008) and the non-targeting shNS is derived from that described in (Tang et al., 2016).

### SA-βgal staining

SA-βgal staining was done using SPIDER-βgal senescence detection kit (Dojindo) according to the manufacturer’s recommendations for fixed cells.

### RNAi-mediated depletion

siRNAs against HP1α (CBX5) Hs_CBX5_5, Cat. No. SI03146479 and All Stars negative control SI03650318 siRNAs were from Qiagen. siRNA was transfected with lipofectamine RNAiMAX (Invitrogen) according to the manufacturer’s protocol. Experiments were performed 36 to 48 hrs post-transfection with individual siRNAs. Depletion was verified in each transfection by Western blotting.

### Antibodies

Antibodies were as follows: rabbit α-biotin Cat. No. A150-109A (Bethyl); mouse α-BrdU/CldU Cat. No. NBP2-44055 (Novus); rabbit α-STING Cat. No.19851-1-AP (Proteintech), mouse α-NCL Cat. No. 396400 (Life Technologies), mouse α-LaminA/C Cat. No. sc-376248 (Santa Cruz Biotechnology), rabbit α-GAPDH Cat. No. 5174 (CST), rabbit α-Histone H3K9me3 Cat. No. 13969 (CST), rabbit α-Histone H3K9me3 Cat. No. A2217P (Diagenode), mouse α-WRN, Cat. No. W0393 (Millipore-Sigma), rabbit α-WRN, Cat. No. NB100-471 (Novus); mouse α-HP1α Cat. No. NBP2-52420 (Novus), rabbit α-HP1α Cat. No. 2616 (CST); rabbit α-V5 Cat. No. 3202 (CST); rabbit α-KAP1 Cat. No. 4124 (CST); mouse α-HDAC2 Cat. No. 5113 (CST); rabbit α-HDAC2 Cat. No. 57156 (CST); rabbit α-LBR Cat. No. A5468 (ABclonal); mouse α-Lamin B1 Cat. No. 66095-1 (ProteinTech).

### Western blotting

Proteins were visualized on Western blots by ECL (Thermo Scientific) and quantified using FluorChem Imager (Alpha Inotech). For presentation, images were saved in TIFF format, adjusted for brightness/contrast and cropped in GIMP, then assembled into figures in CorelDraw. Image brightness/contrast adjustments were made across all lanes of each protein measured. In some cases, lane order was changed and extra lanes were deleted.

### Chromatin immunoprecipitation (ChIP)

ChIP was performed according to the Abcam protocol as described in (Shukla et al., 2022), with 5μg of chromatin using rabbit α-Histone H3K9me3 Cat. No. A2217P (Diagenode) antibody or the equivalent amount of rabbit IgG Cat. No. C15410206 (Diagenode). Input and pulldown DNA was analyzed in triplicates by qPCR with iTaq Universal SYBR Green supermix (Bio-Rad) and the following pairs of primers:

LINE1 5’UTR: 5’GATGATGGTGATGTACAGATGGG3’ and 5’AGCCTAACTGGGAGGCACCC3’ (Shen et al., 2021)

SATII: 5’TCATCGAATGGAAATGAAAGGAGTCATCATCT3’ and 5’CGACCATTGGATGATTGCAGTCAA3’ (Shukla et al., 2022);

αSAT (CST Cat. No. 4486). Only the experiments in which IgG pulldowns were uniformly low (less than 1/10 of the value for H3K9me3) were used for further analysis.

### RNA isolation and RT qPCR

RNAs were isolated using RNeasy Plus RNA isolation kit (Qiagen) and additionally purified of DNA contamination using DNA-free kit (Thermo Fisher). 2μg of RNA was reverse-transcribed using High Capacity cDNA Reverse Transcription kit (Applied Biosystems) per manufacturer’s protocol, with GAPDH-reverse primer 5’GATGCAGGGATGATGTTCTG3’ and HSATII primer 5’ATCGAATGGAAATGAAAGGAGTCA3’. No RT controls were performed for every sample. cDNAs were diluted 1:2 and 1μl of diluted cDNA was used per qPCR reaction. Reactions were run in triplicates with iTaq Universal SYBR Green supermix (Bio-Rad) and the following pairs of primers: 5’ACAACTTTGGCATTGAA3’ and 5’GATGCAGGGATGATGTTCTG3’ for GAPDH; 5’TCATCGAATGGAAATGAAAGGAGTCATCATCT3’ and 5’CGACCATTGGATGATTGCAGTCAA3’ for HSATII. After verifying that no-RT negative control reactions displayed higher Ct values than +RT reactions, the data were processed as follows. Triplicate Ct values for GAPDH were averaged and subtracted from HSAT values to derive ΔCt values, which in turn were used to derive ΔΔCt values for differences between samples within each experiment, and the fold change values (as 2^-ΔΔCt^). Significance was determined in paired two-tailed t-tests performed on ΔΔCt values.

For LMNB1 and LBR RT qPCR, the procedures were as above except that RT reactions were performed with random primers. *LMNB1* and *LBR* (exonic) primers used are described in (Wazir et al., 2013), and the intronic LBR primer is 5’ CACACATCTTTCTGGCGG 3’.

### Proximity Ligation Assay (PLA) and Immunofluorescence (IF) in situ

For PLA detection of protein proximity to nascent DNA, cells were labeled with 20μM EdU (Click Chemistry Tools) for 20 min and EdU was clicked to a mixture of Alexa488 and biotin azides at molar ratio of 1 to 50. Protein/protein PLA was performed according to the standard procedures. PLA DuoLink red or green detection reagents and DuoLink anti-mouse and anti-rabbit antibodies (Millipore-Sigma Cat. No DUO92008, DUO92014, DUO92001, and DUO92002, respectively) were used as described previously (Lazarchuk et al., 2019), except that after formaldehyde fixation cells were washed in PBS, permeabilized by addition of 4°C 90% methanol in PBS and stored at −20°C prior to staining. The staining was preceded by two washes in PBS, an incubation with 0.25% triton X100 for 10 min, and two more washes in PBS. IF is situ was performed as described before (Lazarchuk et al., 2020) except in the case of KAP1 IF, which was performed according to the manufacturer’s protocol. Images of cells were collected under Zeiss Axiovert 200M microscope with 40X magnification objective using Micro Manager software. Digital images were analyzed with Fiji ImageJ software package with custom macros as described in (Lazarchuk et al., 2019) or with the Cell Profiler software package.

### Fiber FISH and microfluidics assisted replication track analysis (maRTA)

DNA stretching was performed as described before (Kehrli et al., 2016; Sidorova et al., 2009). After drying at room temperature overnight, coverslips were incubated in 2XSSC, 70% formamide at 75C for 5 min, dehydrated in 70, 85, 95, 100% ice-cold ethanol series for 2 min each, and air-dried. Hydridization mixture containing 50% formamide, 20mM Tris HCl pH7.4, 5mg/ml blocking reagent (Roche), and 0.5ug/ml biotinylated LNA probe against SATII (based on the sequence ATTCCATTCAGATTCCATTCGATC (Swanson et al., 2013) and custom-produced by IDT) was incubated at 80C for 5 min and aliquoted onto a glass slide prewarmed to 80C. Coverslips were placed DNA side down into hybridization mixture droplets, incubated at 80C for 5 min and then placed into a humidified chamber at room temperature for 2 hrs. Coverslips were then washed in 2xSSC, 01% Tween 20, once at room temperature and once at 50C, for 10 min each. Coverslips were rinsed in maRTA antibody wash buffer, then blocked and stained with rabbit α-biotin Cat. No. A150-109A (Bethyl); mouse α-BrdU/CldU Cat. No. NBP2-44055 (Novus) antibodies followed by secondary antibodies as in (Sidorova et al., 2009). Microscopy of stretched DNAs was performed on the Zeiss Axiovert microscope with a 40x objective as above. Lengths of tracks were measured in raw merged images using Zeiss AxioVision software. Fluorochromes were Alexa594 for CldU, Alexa488 for biotin.

### Statistical analysis

Statistical analyses and graphing of the data were done in R studio. P values for qPCR results were derived from pairwise t-tests on ΔΔCq values. P values for the rest of the assays were as follows. For continuous variables (e.g., track lengths, mean fluorescence intensities) p values were calculated in K.S. tests and for discrete variables (e.g. PLA foci) – in Wilcoxon tests. The analyses were always performed on whole datasets, without exclusion of any outlier data points. To quantify and concisely visualize the sizes of differences between PLA foci, track lengths, or MFI distributions of pairs of samples we calculated the Cliff’s delta statistic for these pairs of samples. Cliff’s delta is a metric recommended for comparison of non-parametric distributions. In general, Cliff’s delta of a distribution A vs. distribution B can range from 1 (if all values in A are larger than all values in B) to −1 (if the reverse is true), and 0 value indicates that the distributions are completely overlapping. For fiber FISH/maRTA, p value is the probability with which a subset the size of n = n of C_SAT_ and a mean value equal to that of C_SAT_ can be randomly drawn from a full CldU track dataset (C_gen_+ C_SAT_). To calculate it, the random subset of n = n of C_SAT_ was drawn 100 times from the full set, the means of each subset were plotted in a frequency distribution, and a two-sided one-sample t test was performed on this distribution for the null hypothesis that its mean equals the mean of C_SAT_ at a confidence level= 0.98.

## Results

### WRN associates with HDAC2

We developed PLA assays to visualize WRN/HDAC2 and WRN/HDAC1 physical proximities *in situ*. Robust WRN/HDAC2 PLA signal was detected in the HDAC2+ GM639 cell line derivative and was dramatically reduced in the isogenic hdac2-27 null cells (Fig 1A). A V5-tagged version of HDAC2 was introduced into hdac2-27 cells (Fig. 1B,C), and a V5/WRN PLA signal was observed in these cells both in S phase and outside it, as identified by pulse-labeling with EdU (Fig. 1D). Thus, the WRN/HDAC2 association was not limited to nascent chromatin. A moderate increase of the signal in EdU+ compared to EdU-cells can be attributed to an increase in chromatin content per nucleus. WRN-dependence of the WRN/HDAC2 PLA signal was also confirmed with a derivative of WI38hTERT fibroblasts in which WRN was stably depleted by doxycycline-inducible shRNA expressed from a lentiviral vector (FIG.1,E,F; unless stated otherwise, here and elsewhere WI38hTERT cells were maintained in a WRN-depleted state for a number of passages to approximate a steady state of WRN deficiency).

**Figure 1.**
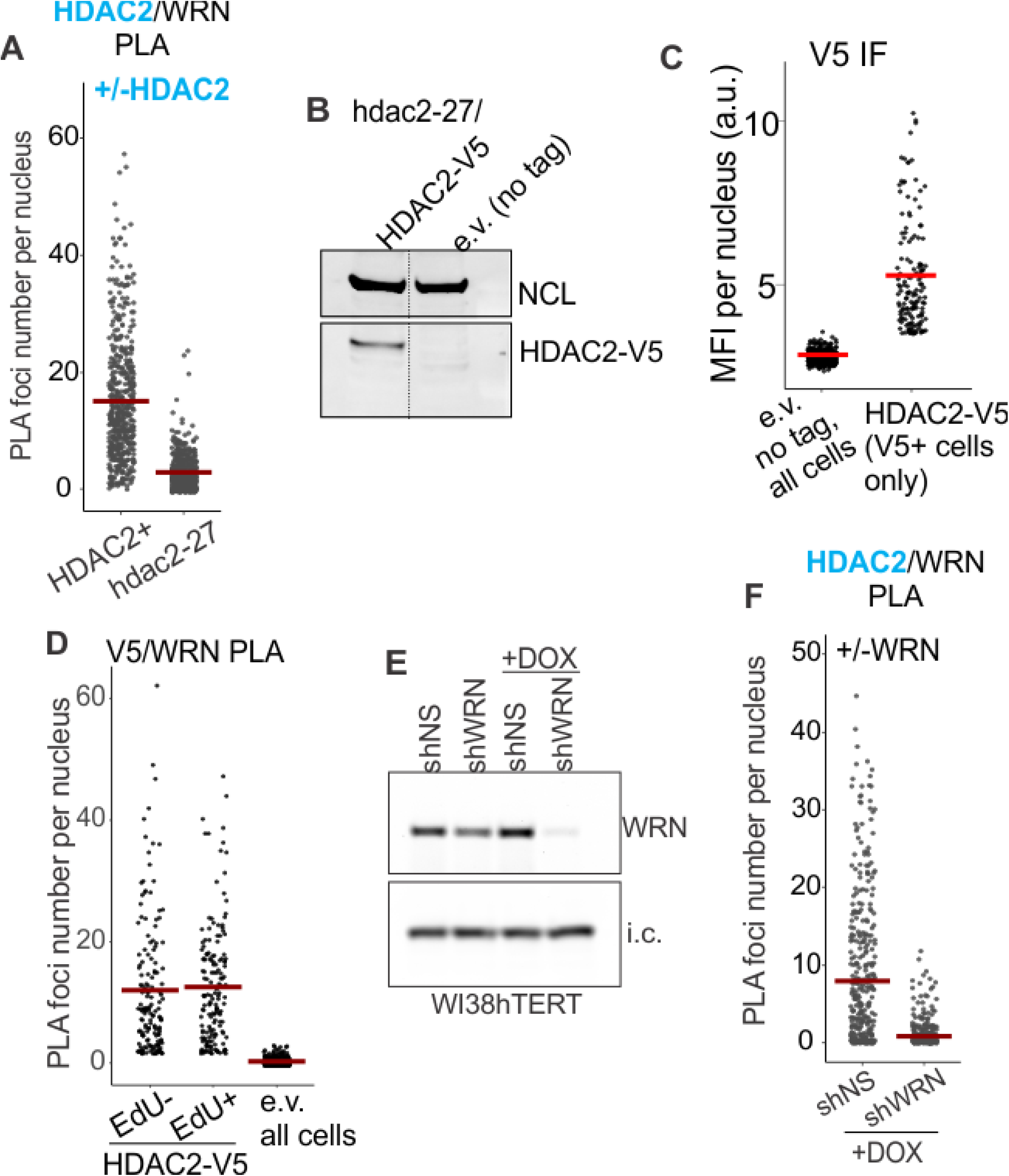
WRN associates with HDAC2. A) Quantitation of HDAC2/WRN PLA performed with the isogenic HDAC2+ and hdac2-27 null derivatives of GM639. B) Hdac2-27 cells were stably transfected with HDAC-V5 transgene or empty vector (tag-less) and analyzed by Western blotting with antibodies to NCL (internal control) and V5. C) Quantitation of IF in situ performed with V5 antibody on the same cells as in (B). MFI, mean nuclear fluorescence, a.u., arbitrary units. D) Quantitation of V5/WRN PLA performed on the same cells as in (B,C), labeled for 30 min with EdU. EdU incorporation was visualized by Clicking to Alexa 488 azide. V5/WRN signals of EdU-positive and EdU-negative cells are plotted separately. E) WI38hTERT cells were stably transfected with the indicated shRNA-expressing constructs and treated with 100ng/ml doxycycline for 5 days prior to Western blotting with WRN antibody. i.c., internal control (the STING protein). F) Quantitation of HDAC2/WRN PLA performed with the same cells as in (E). Crossbars in jitter plots are distribution means.

To test the contribution of HDAC1, we used clonally derived control and hdac1 null normal human dermal fibroblasts (NHDFs) in which HDAC1 was disrupted (FIG.S1A) using CRISPR/Cas9 as described before (Lazarchuk et al., 2020). We detected a more robust signal of PLA foci of WRN/HDAC2 compared to WRN/HDAC1, the latter measuring virtually at a background level (FIG.S1B, C). The WRN/HDAC2 signal was once again present in cells outside of S phase and was not reduced in hdac1 null cells (FIG.S1C), suggesting that the WRN/HDAC2 association does not require HDAC1. A mild elevation of the HDAC2/WRN signal in hdac1 null compared to the HDAC1 wt control is consistent with the compensatory increase in HDAC2 levels upon HDAC1 disruption, as we and others observed before (Lazarchuk et al., 2020; Yamaguchi et al., 2010). Together, these observations led us to focus on the WRN/HDAC2 association for further study.

### WRN and HDAC2 associate with heterochromatin protein 1 alpha (HP1α) independently of each other and depend on it for mutual proximity

Given that WRN/HDAC2 association was not limited to S phase cells, we reasoned that the putative function of this association is not centered on replication forks or nascent chromatin. Previous studies suggest an alternative substrate for potential physical and functional interaction of WRN and HDAC2: CH, particularly pericentromeric heterochromatin, PCH (Craig et al., 2003; Vaute et al., 2002; Zhang et al., 2015). WRN and HDAC2 were found to co-immunoprecipitate with heterochromatin protein 1 alpha, HP1α, a major component of CH and PCH (Fioriniello et al., 2020; Meyer-Nava et al., 2020), albeit the functional significance of these interactions is not yet known.

To confirm WRN/HP1a and HDAC2/ HP1α associations, we used the hdac2-27 null and complemented cells described in the previous section, a CRISPR/Cas9 WRN KO derivative of GM639 Δwrn-3 (Fig.2A) that was stably transfected with a WRN-expressing construct (Fig.2B), the shRNA-mediated depletion of WRN described in the previous section (FIG.1), and siRNA-mediated depletion of HP1α (FIG.2C,D). Pairwise associations between HDAC2, HP1α, and WRN in cell nuclei were assessed by PLA in situ.

**Figure 2.**
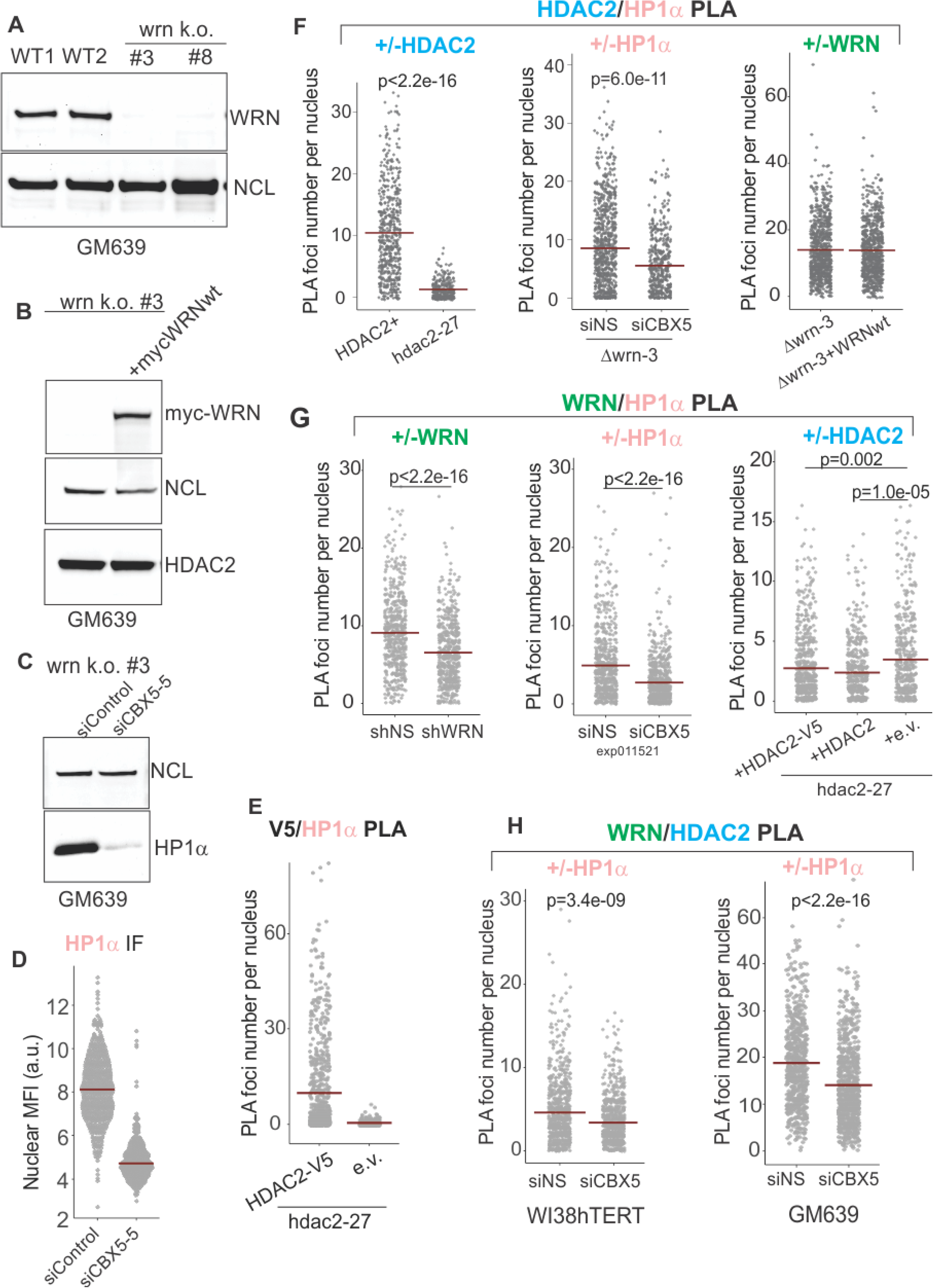
WRN and HDAC2 proximity is facilitated by HP1α. A) Clonal isolates of CRISPR/Cas9 knockouts of WRN were generated in the GM639 background. WT1 and WT2 are isogenic WRN wild type controls. WT1 (GM639-Cas9-EV (Lazarchuk et al., 2020)), parent cell line is a GM639 derivative with Cas9 and an empty vector instead of a dgRNA expressing construct. WT2 is one of WRN wild type isolates derived from subcloning of WRN dgRNA-transfected GM639-Cas9 cells. B) Western blot analysis of the *WRN* k.o. #3 cell line (hereafter called Δwrn-3) stably transfected with pLX209-neo-WRN expressing 5xmyc-WRN. C) Δwrn-3 was transfected with non-targeting (siControl) and the *HP1α* gene-targeting (CBX5-5) siRNAs and analyzed by Western blotting at 36 hrs post-transfection. D) WI38hTERT expressing non-targeting shNS were transfected with non-targeting and *HP1α*-targeting (CBX5-5) siRNAs and analyzed by IF in situ with HP1α antibody at 36 hrs post-transfection. E) Quantitation of V5/HDAC2 PLA performed in hdac2-27 cells stably transfected with HDAC2-V5 transgene or with empty vector (e.v., no tag). Note that not all transfected cells express HDAC2-V5, thus cells with no PLA signal are expected in the hdac2-27/HDAC2-V5 population. F) Quantitations of HDAC2/HP1a PLA performed in Δwrn-3 and hdac2-27 (left panel), in Δwrn-3 transfected with non-targeting or CBX5-5 siRNAs (center panel), or in Δwrn-3 with or without expression of the 5xmyc-WRN (right panel). The graph represents two independent experiments. G) Quantitations of WRN/HP1a PLA performed in WI38hTERT expressing non-targeting shRNA or shRNA against WRN (left panel), in WI38hTERT expressing non-targeting shRNA and also transfected with siRNA against HP1α or a non-targeting control (center panel, the experiment was done at 36 hrs post-transfection); and in hdac2-27 stably expressing the indicated transgenes or an empty vector (right panel, the graph represents three independent experiments). H) Quantitations of WRN/HDAC2 PLA performed in WI38hTERT with siRNA against HP1α or a non-targeting control (left panel) and in WT1 (GM639-Cas9-EV) transfected with the same siRNAs (right panel). P values were derived in Wilcoxon tests. Crossbars in graphs are distribution means.

We detected a HDAC2-V5 transgene-dependent V5/HP1a PLA signal (FIG.2E) as well as a HDAC2/HP1α PLA signal that was responsive to HDAC2 and HP1a manipulation (FIG.2F, left and center panels) but was unaffected by WRN manipulation (FIG.2E, right panel; also see FIG.4F, left panel). Likewise, a WRN and HP1α-dependent PLA signal was readily detectable (FIG.2F left and center panels), and if anything was modestly increased in the absence of HDAC2 (FIG.2F, right panel). Remarkably, depletion of HP1α in WI38hTERT or GM639 cell backgrounds also reduced the HDAC2 to WRN PLA signal (FIG.2G). These data suggest that HP1a provides a physical platform for, or regulates the assembly of a higher order complex within which WRN and HDAC2 are brought into proximity. At the same time, WRN and HDAC2 do not require each other to associate with HP1α.

### Senescing cells downregulate WRN, HDAC2, HP1α associations

Given the findings that HDAC2 (Wagner et al., 2001) and WRN (Zhang et al., 2015) complexes may be altered in senescing cells, we examined how two distinct types of senescence, replicative (RS) and oncogene-induced (OIS) affect the state of WRN, HDAC2, HP1α associations. NHDFs were serially passaged and cryopreserved at various times to generate early and late population doubling level (PDL) cultures to compare side by side later. The PDL5 (“young”) and PDL47 (“old”) fibroblasts were chosen for comparison (FIG.3A, PDL values correspond to those at the beginning of the experimental series). The old fibroblasts exhibited slower growth and larger size but were not yet fully senesced, presenting a heterogeneous population of cells with normal or enlarged nuclei and an overall elevated level of senescence-associated β-gal staining (FIG.3B). Old fibroblasts showed reductions in mean fluorescence intensity of H3K9me3, HDAC2, HP1α, and WRN compared to proliferating young cells in IF in situ assays (FIG.3C,D). The levels of these proteins in whole cell extracts were comparable in old and quiescent young fibroblasts and virtually the same or slightly higher in proliferating young fibroblasts (FIG.3E). Nevertheless, all three of pairwise associations, HP1α/WRN, HDAC2/WRN, and HDAC2/ HP1α, were reduced in old fibroblasts compared to young (FIG.3F and FIG.S2), consistent with the reduced mean nuclear levels of the individual partners to these associations.

**Figure 3.**
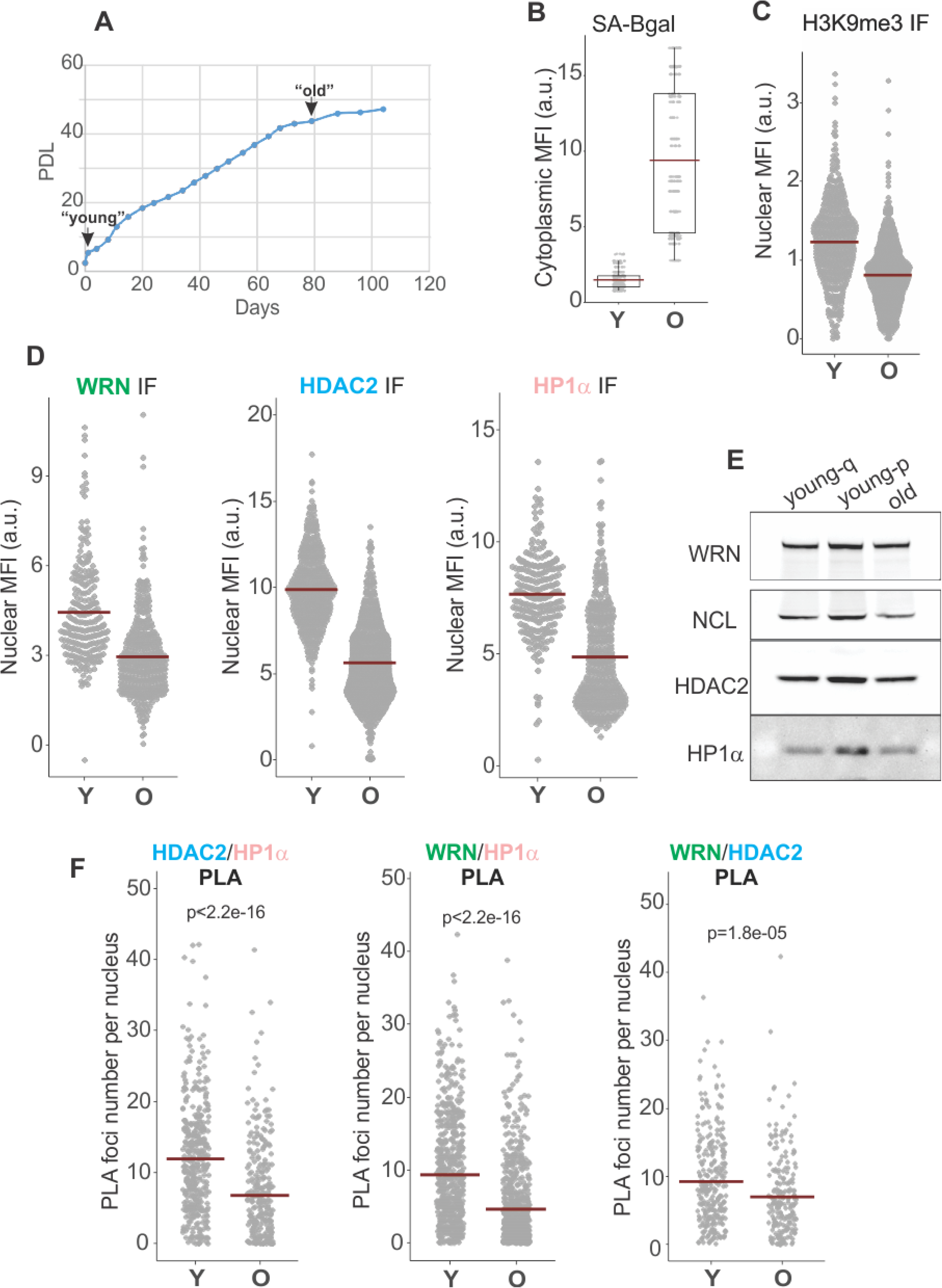
WRN, HDAC2, and HP1a associations are reduced in replicatively senescing cells. A) Normal human dermal fibroblasts were passaged to derive early and late passage cultures, which were cryopreserved and subsequently used in the same experiment. B) SA-βgal levels in young (Y) and old (O) fibroblasts were quantified via fluorescent detection and microscopy. Cytoplasmic MFI was measured per digital image, normalized to the number of cells in each image determined by DAPI counterstaining, and plotted. C,D) Quantitation of H3K9me3, WRN, HDAC2, and HP1a mean nuclear levels in young vs. old fibroblasts determined by IF in situ. E) Total levels of the same proteins in old vs. quiescent (young-q) and proliferating (young-p) young fibroblasts determined by Western blotting. F) Quantitations of HDAC2/HP1a, WRN/HP1a, and WRN/HDAC2 PLAs in young vs. old fibroblasts. Graphs in C, D, and F represent two independent experiments each. P values were derived in KS tests for MFI values and Wilcoxon tests for PLA foci values. Crossbars in graphs are distribution means.

Oncogene-induced senescence, OIS, triggered by the overexpression of oncogenes such as RAF or RAS shares some features with replicative senescence, some of the key differences being a rapid, abrupt onset and the spatial reorganization of chromatin into senescence-associated heterochromatin foci, or SAHF (Narita et al., 2003), which appear to reflect contraction of the chromosomal territories (Chandra et al., 2015; Chandra et al., 2012; Chandra and Narita, 2013; Lenain et al., 2015; Sadaie et al., 2013) (FIG.4A). This, however, does not indicate a global heterochromatin dismantlement in RAF OIS (Hao et al., 2022; Sati et al., 2020; Sun et al., 2018). At the same time, CH domains, including pericentromeres, may be selectively relaxed in OIS (Dillinger et al., 2017; Mendez-Bermudez et al., 2022; Swanson et al., 2013). We used an inducible RAF model system (Jeanblanc et al., 2012) to trigger OIS in WI38hTERT by addition of 4-HT to the cell media (FIG.4A). Nuclear MFI of H3K9me3 was consistently higher in OIS cells compared to proliferating controls (FIG.4B). Likewise, HP1α, HDAC2, and WRN nuclear MFI were the same or higher than the non-OIS controls (FIG.4C). However, combined with the fact that the levels of HDAC2, WRN, and HP1α in whole cell extracts from OIS cells were comparable to those in proliferating cells (FIG.4D), the MFI increases could at least in part be explained by a reduction of nuclear area in OIS. Strikingly however, the HP1α /WRN, HP1α /HDAC2, and WRN/HDAC2 PLA signals were drastically downregulated (FIG.4E, F and FIG.S3). Taken together, the data suggest that reduction of associations between these proteins may be linked to a subset of heterochromatin changes that are shared between replicative senescence and OIS, which could be the relaxation of CH.

**Figure 4.**
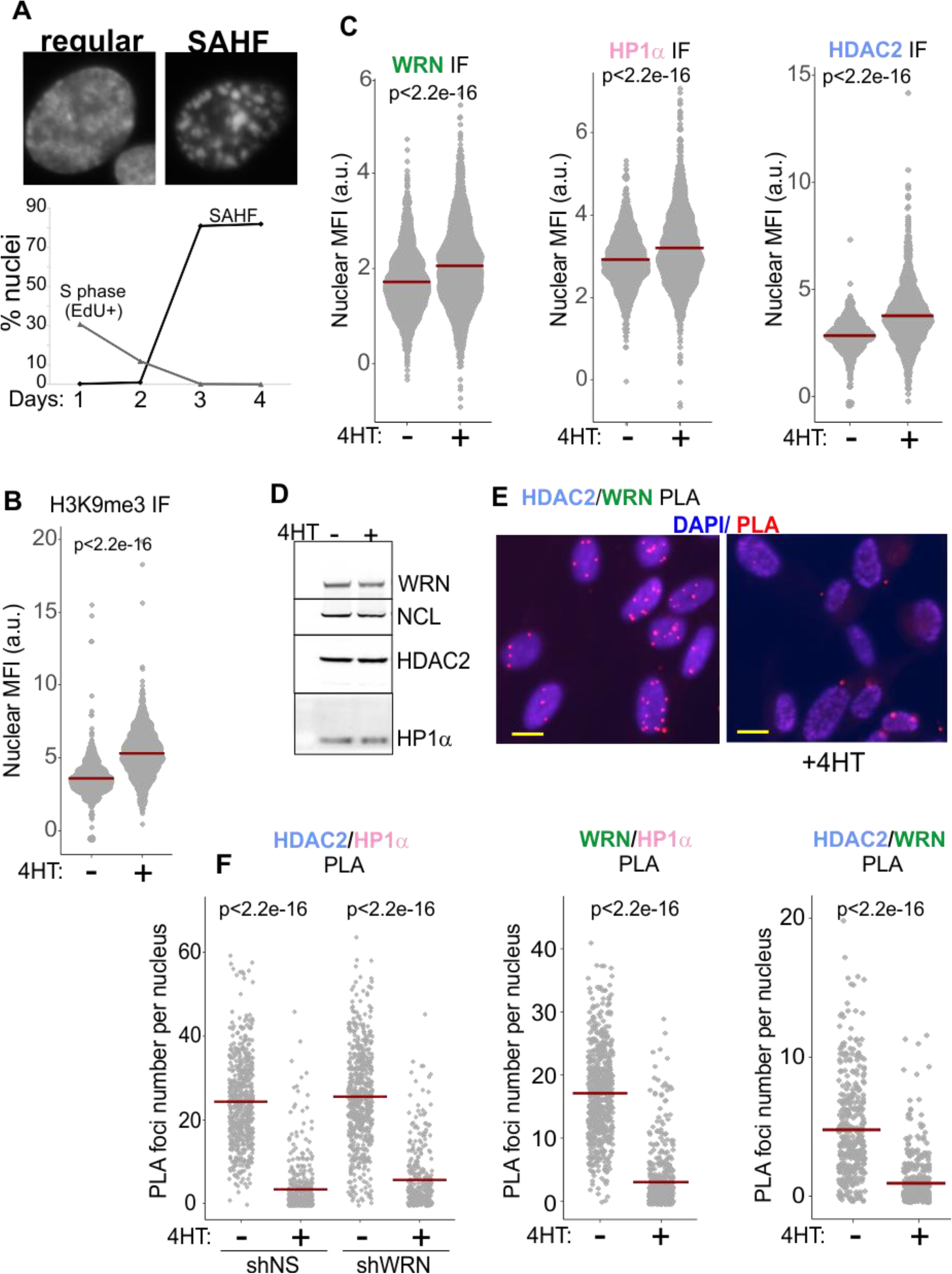
WRN, HDAC2, and HP1a associations are reduced in RAF oncogene-induced senescence. A) 4-HT-inducible expression of RAF oncogene in WI38hTERT leads to cessation of proliferation and onset of senescence marked with formation of SAHF after three days. B-C) Quantitations of H3K9me3 (B, three independent experiments), and WRN, HP1a, and HDAC2 (C, two independent experiments each) nuclear levels in WI38hTERT after 4 days in 4-HT compared to the contemporaneous control without 4-HT, determined by IF in situ. D) Total levels of the WRN, HP1a, and HDAC2 proteins in WI38hTERT under the same conditions, determined by Western blotting. E) An example of HDAC2/WRN PLA data in WI38hTERT with and without 4HT. Scale, 10um. F) Quantitations of HDAC2/HP1a, WRN/HP1a, and HDAC2/WRN PLA in WI38hTERT after 4 days in 4-HT compared to the contemporaneous no-4HT control. The left and center panels represent two independent experiments each, and the right panel represents two biological replicas. P values were derived in KS tests for MFI values and Wilcoxon tests for PLA foci values. Crossbars in graphs are distribution means.

### Constitutive heterochromatin maintenance in WRN-deficient proliferating cells

We next asked if WRN deficiency is sufficient to elicit CH defects in proliferating, immortalized cells. While the overall H3K9me3 signal intensity in situ was somewhat increased in WRN-depleted WI38hTERT compared to the isogenic control (FIG.5A), there was a decrease of H3K9me3 on SATII and αSAT classes of satellite repeats comprising PCH but, interestingly, not on the LINE1 interspersed repeats representing other CH (FIG.5B). To follow this up, we looked at SATII expression levels in cells with and without WRN. In WI38hTERT, SATII transcripts were difficult to detect however (FIG.S4A, left panel), unless the cells were induced to enter OIS, where, as expected (reviewed in (Rocha et al., 2021)), the level of SATII transcripts was significantly upregulated (FIG.S4A, right panel). Though WRN-depleted WI38hTERT showed somewhat higher SATII RNA levels both in proliferating and OIS cells, the data were not robust enough to reach significance (FIG.S4A). In contrast, we were able to robustly measure SATII RNA in proliferating GM639, where complementation of *wrn* null cells with the wild type WRN led to a marked reduction in SATII RNA level (FIG.5C), suggesting that silencing of SATII repeats was compromised in *wrn* null cells.

**Figure 5.**
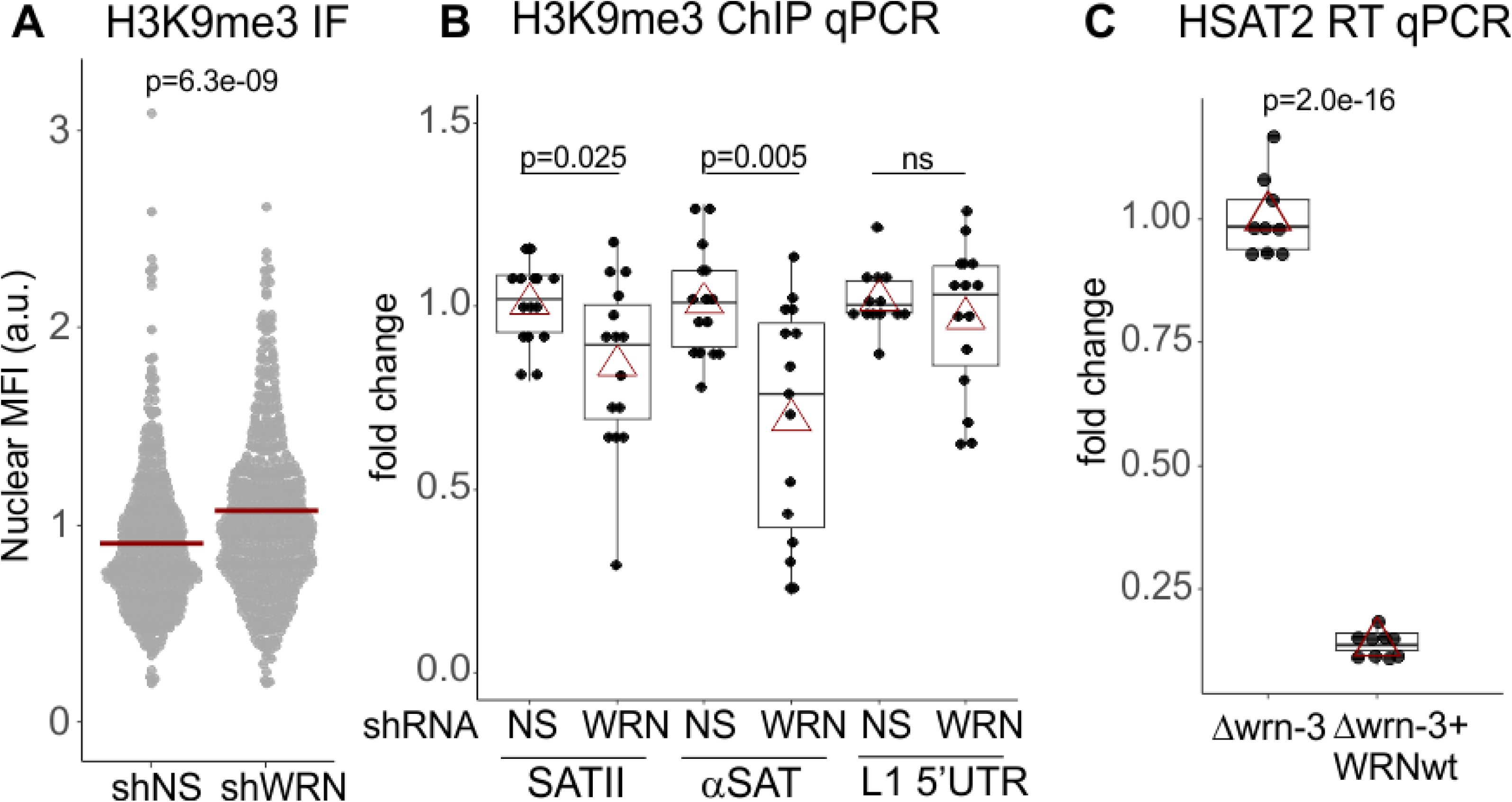
H3K9me3 heterochromatic mark is reduced on satellite tandem repeats in WRN-depleted cells. A) Quantitation of IF in situ for H3K9me3 levels in WI38hTERT expressing shRNA against WRN or non-specific control (NS). Crossbars are distribution means, and the p value was determined by a KS test. The graph represents three independent experiments. B) ChIP analyses of H3K9me3 levels on the indicated genomic loci, performed in the same cells as in (A). The graph summarizes results of five independent experiments with technical triplicates. C) RT qPCR analyses of SATII satellite repeat transcription performed on Δwrn-3 cells with and without complementation by the wild type WRN gene. The graph summarizes three independent experiments with technical triplicates. In (B) and (C), red triangles are means and boxplots mark first and fourth quartiles and the distributions’ medians. P values were calculated in paired t-tests carried out on ΔΔ Cq values.

CH comprises a network of HP1α interactions, including with Lamins and with Lamin receptors anchored in the inner nuclear membrane (Bizhanova and Kaufman, 2021). The latter interactions participate in tethering of heterochromatin to the nuclear periphery (Chang et al., 2022). We hypothesized that CH maintenance in WRN-deficient cells may be affected by means of altering other relevant interactions of HP1α, and examined its well-known associations with KAP1 and the Lamin B receptor LBR (Kumar and Kono, 2020; Meyer-Nava et al., 2020), which are important for heterochromatin assembly (FIG.6, see FIG.S5A for PLA controls).

**Figure 6.**
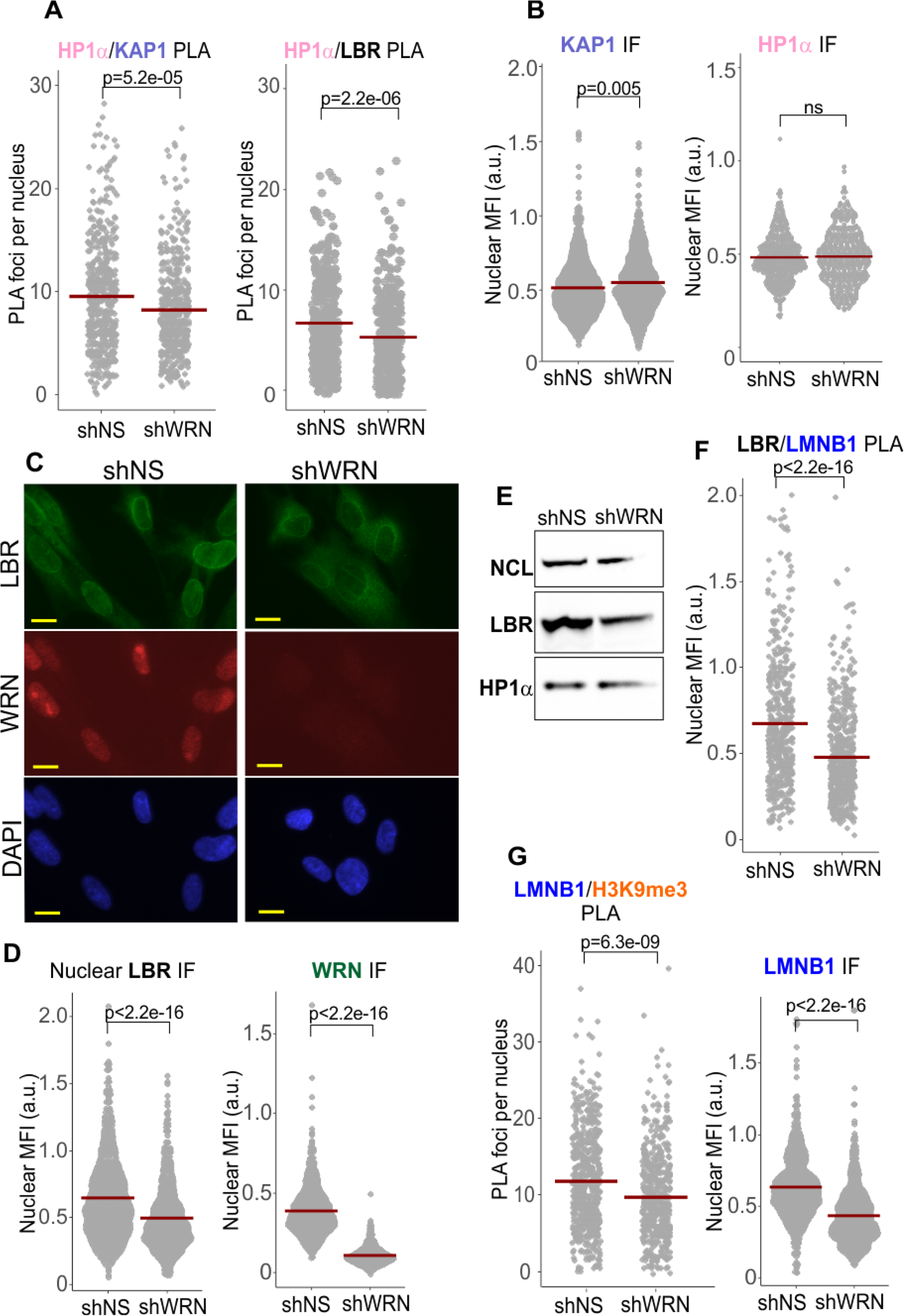
Reduction of Lamin B1-associated interactions in WRN-depleted cells. A, B) Quantitations of the indicated PLA and IF in situ analyses in WI38hTERT cells expressing shRNA against WRN or control shRNA. Graphs in (A) represent two (HP1a/KAP1), four (HP1a/LBR), and in (B) two (KAP1), and three (HP1a) independent experiments each. C) An example of LBR and WRN IF in situ in WI38hTERT cells expressing the indicated shRNAs. Scale, 10μm. D) Quantitations of the indicated IF in situ analyses in the same WI38hTERT cells as in (C). The panels represent over five (WRN) and four (LBR) independent experiments each. E) A Western blot of WI38hTERT expressing the indicated shRNAs and probed with antibodies against NCL (internal control), LBR, and HP1α. F, G) Quantitations of the indicated PLA or IF in situ analyses in the same WI38hTERT cells above. The panels represent, two (LBR/Lamin B1), three (Lamin B1/H3K9me3), and three (Lamin B1) independent experiments each. Nuclear MFI values were used instead of PLA foci numbers in cases where the latter numbers were too high to robustly count individual foci. P values were derived in Wilcoxon tests for PLA foci values and KS tests for the MFI values.

HP1α association with KAP1 was modestly reduced in WRN-depleted WI38hTERT (FIG.6A, left panel). KAP1 nuclear level measured by IF in situ, was, if anything, slightly increased by WRN depletion (FIG.6B, left panel). HP1α nuclear level trended as largely unchanged upon WRN depletion (FIG.6B, right panel), though one of the four independent measurements demonstrated a WRN-dependent reduction (FIG.S5B). It is possible that this variability reflects the complexity of intra-and intermolecular interactions of HP1α, which can affect the accessibility of its epitope *in situ*. No difference in HP1α levels was detected in whole cell extracts of WRN-depleted and control cells by Western blotting (FIG.6E).

HP1α association with LBR trended toward reduction in WRN deficient cells (FIG.6A, right panel, and FIG.S5B). LBR, a transmembrane protein, can localize to the INM and ER, with the INM typically being its predominant location (Nikolakaki et al., 2017). Interestingly, in WRN-deficient WI38hTERT LBR appeared comparatively less enriched within the perimeter of the nucleus (FIG.6C and FIG.S5C, hereafter called nuclear LBR). Since quantitation of the nuclear signal intensity involves normalization to the fluorescence background outside the nuclei, for LBR this metric incorporates its enrichment in the nucleus vs. the cytoplasm. Indeed, in WRN-deficient cells LBR localization pattern translated into a significant reduction of the nuclear LBR signal (FIG.6D). In addition, the LBR protein level was reduced in whole cell extracts of WRN-depleted cells on Western blots (FIG.6E). Consistent with a relative reduction of the nuclear LBR, Lamin B1/LBR proximity was also significantly reduced without WRN (FIG.6F), which was also accompanied by a reduced nuclear level of Lamin B1 (FIG.6G, right panel), and a reduced Lamin B1/H3K9me3 proximity (FIG.6G, left panel).

Marked reduction of the nuclear LBR and LMNB1/LBR PLA signals was also seen in WI38hTERT newly depleted of WRN (at day 5 post induction of shRNA, FIG.S4B, C). Lamin B1 was moderately downregulated and the H3K9me3 mark upregulated in these cells (FIG.S4B). Both in chronically and newly WRN-depleted cells, lower LBR levels in the nuclei were not associated with a morphologically distinct subset of cells, as can be seen for example from binning LBR levels by nuclear area (FIG.S6A). Rather, these values were lower for virtually every nuclear area bin including those containing the bulk of cells in both WRN-depleted and control populations. In addition, nuclear levels of LBR were lower both in S phase and non-S phase WRN-depleted cells (Fig. S6B). These data argue against a notion that the lower population mean of LBR levels in the WRN-depleted WI38hTERT line is driven by a subset of it that is atypical, e.g. is cell cycle-arrested or senesced.

Nuclear LBR was also lower in Δwrn-3 GM639 fibroblasts compared to the same cells stably transfected with a lentiviral vector-based WRN transgene (FIG.7). The exonuclease or helicase-dead mutants of WRN, respectively, E84A and K577M, transfected and expressed from the same backbone (FIG.7A), were comparable to the wild type WRN in their nuclear LBR levels (FIG.7B). In these experiments, only the LBR fluorescence readouts from WRN-positive cells were included in the measurement since only a fraction of cells in these populations expressed detectable levels of WRN (wild type or mutant, FIG.7A). We could not measure WRN levels and PLA signal simultaneously due to an antibody conflict. With that caveat however, we could still observe that re-expression of the wild type WRN in Δwrn-3 cells led to an increase in Lamin B1/LBR and Lamin B1/H3K9me3 PLA signals compared to the uncomplemented Δwrn-3, whereas the HP1a/KAP1 signal was slightly decreased (FIG.7C). Thus, only the Lamin B1 and LBR associations, levels, and in case of LBR, distribution, were the features consistently reduced in WRN deficiency, whereas HP1α was comparatively less affected.

**Figure 7.**
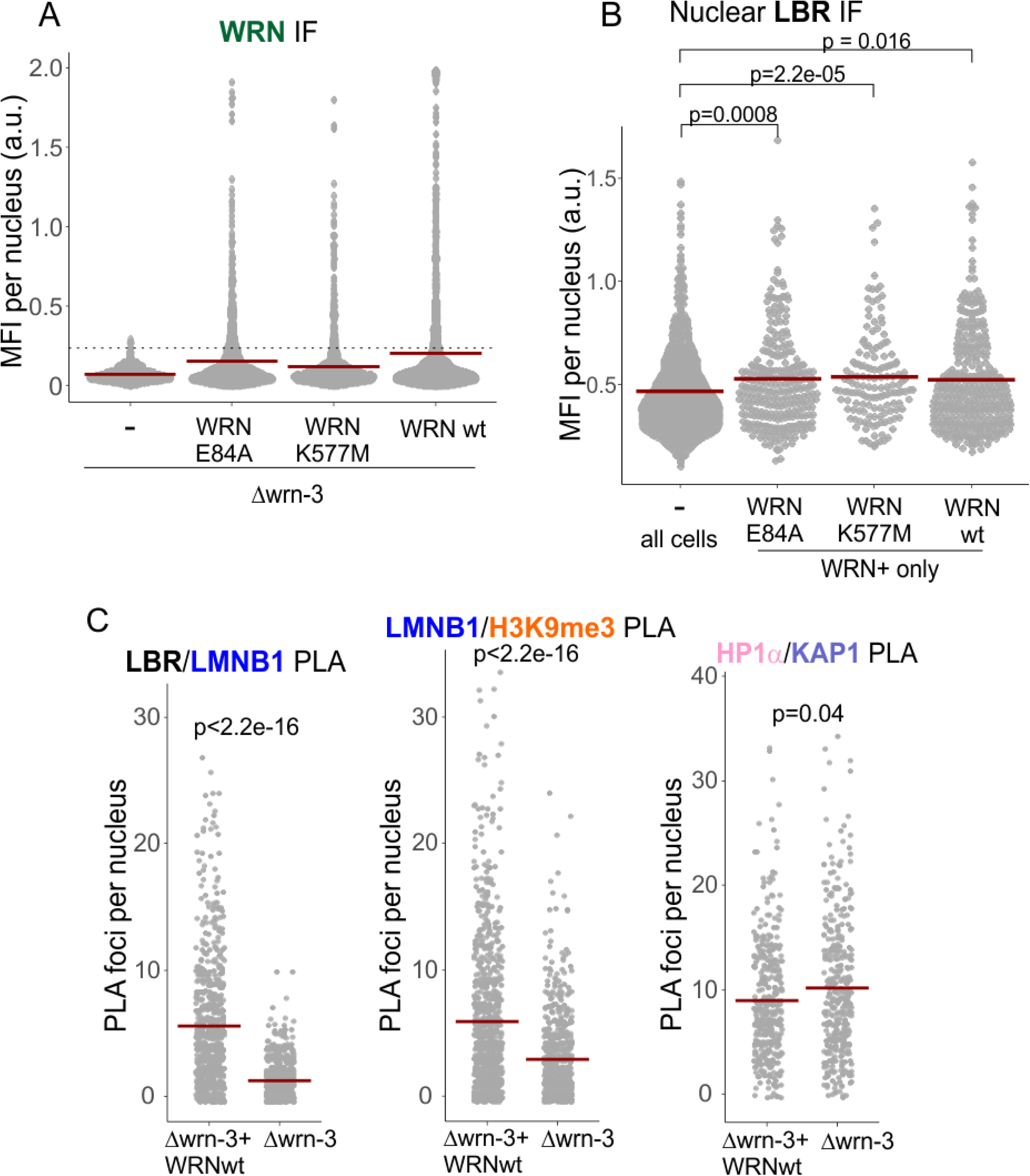
Catalytically inactive WRN mutants restore nuclear LBR levels in *wrn* null cells. A) Stable expression of the wild type, exonuclease-dead (E84A), or helicase-dead (K577M) WRN in Δwrn-3 cells, was measured by IF in situ with an antibody against WRN. A dotted line marks the cutoff MFI below which cells were considered negative for WRN expression. B) Nuclear enrichment of LBR was measured by IF in situ in the same cells as in (A). For WRN-complemented cells, only WRN-positive cells measuring above the cutoff line in (A) for WRN expression were plotted. The graph represents three independent experiments for the wild type and E84A WRN and two independent experiments for the K577M WRN. C) Quantitations of the indicated PLA analyses performed in the Δwrn-3 GM639 with or without expression of the wild type WRN transgene. P values were calculated in KS tests for MFI values and Wilcoxon tests for PLA foci. Crossbars in graphs are distribution means.

Reduction of Lamin B1 levels is a prominent feature of senescence (Dou et al., 2016; Dou et al., 2015; Lenain et al., 2015; Shah et al., 2013; Shimi et al., 2011), and at least in some cell types, Lamin B1 or LBR can trigger senescence when depleted (Herman et al., 2021; Lämmerhirt et al., 2022; Shimi et al., 2011). Indeed, both the nuclear levels of Lamin B1 and LBR (by IF in situ, FIG.8A) as well as associations of Lamin B1/LBR and Lamin B1/H3K9me3 (by PLA, FIG.8B) were also downregulated in replicatively senescing “old” primary, WRN-proficient fibroblasts compared to the isogenic “young” fibroblasts. The HP1α/KAP1 PLA signal was also markedly reduced in old vs. young fibroblasts, while the HP1α/LBR signal was reduced to the least degree of all PLA pairs measured (FIG.8C). Nuclear Lamin B1 level was also profoundly reduced in WI38hTERT undergoing OIS, as expected (FIG. S6C). Interestingly however, cellular levels of LBR did not appear to reduce in OIS cells, rather, subcellular distribution of LBR changed to a prominently perinuclear/cytoplasmic localization (FIG.S6D). Together, the data indicate that WRN-depleted proliferating cells exhibit an overall milder yet overlapping set of perturbations of the heterochromatin complexes as those seen in WRN-proficient senescing cells.

**Figure 8.**
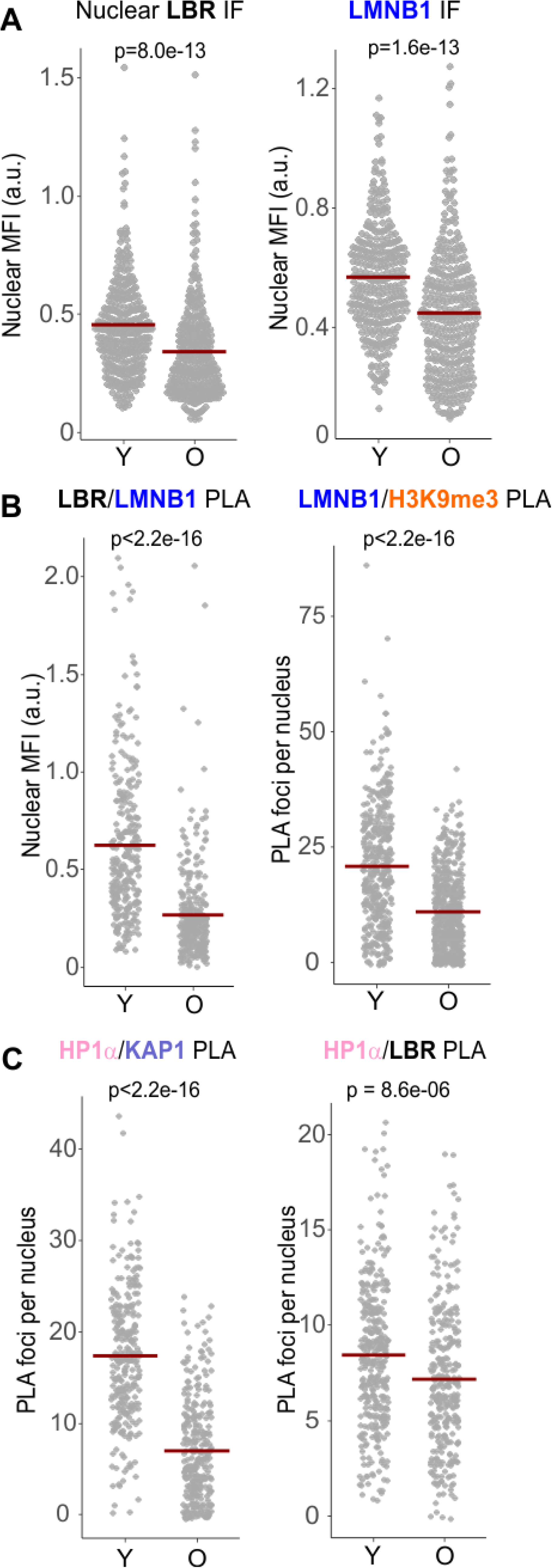
Replicatively senescing cells reduce the levels of LBR, Lamin B1, and the associated protein complexes. A) Quantitations of IF is situ analyses of the levels of LBR and Lamin B1 in the nuclei of early (young, Y) and late passage (old, O) normal human dermal fibroblasts. B,C) Quantitations of the indicated PLA analyses in of the same cells as in (A). All panels represent two independent experiments. P values were calculated in KS tests for MFI values and Wilcoxon tests for PLA foci. Crossbars in graphs are distribution means.

### LBR reduction in WRN-deficient cells is at least in part mediated through transcription

The *LBR* transcript showed up to 50% reduction in WI38hTERT that were WRN-depleted over a long term (FIG.9A), as well as in the newly-depleted WI38hTERT (FIG.S4D). On the other hand, the level of the *LMNB1* transcript was not reduced in WRN-depleted WI38hTERT, and in fact it was slightly elevated upon long-term depletion of WRN (FIG.9A and S4D). Also, LBR transcript was lower in wrn null GM639 cells compared to their wild type WRN-complemented counterparts (FIG.9B). We were not able to reliably compare LBR transcript levels in WRN K577M and E84A mutants vis-a-vis WRN wt because the mutants were expressed in a smaller fraction of cell population than WRN wt.

**Figure 9.**
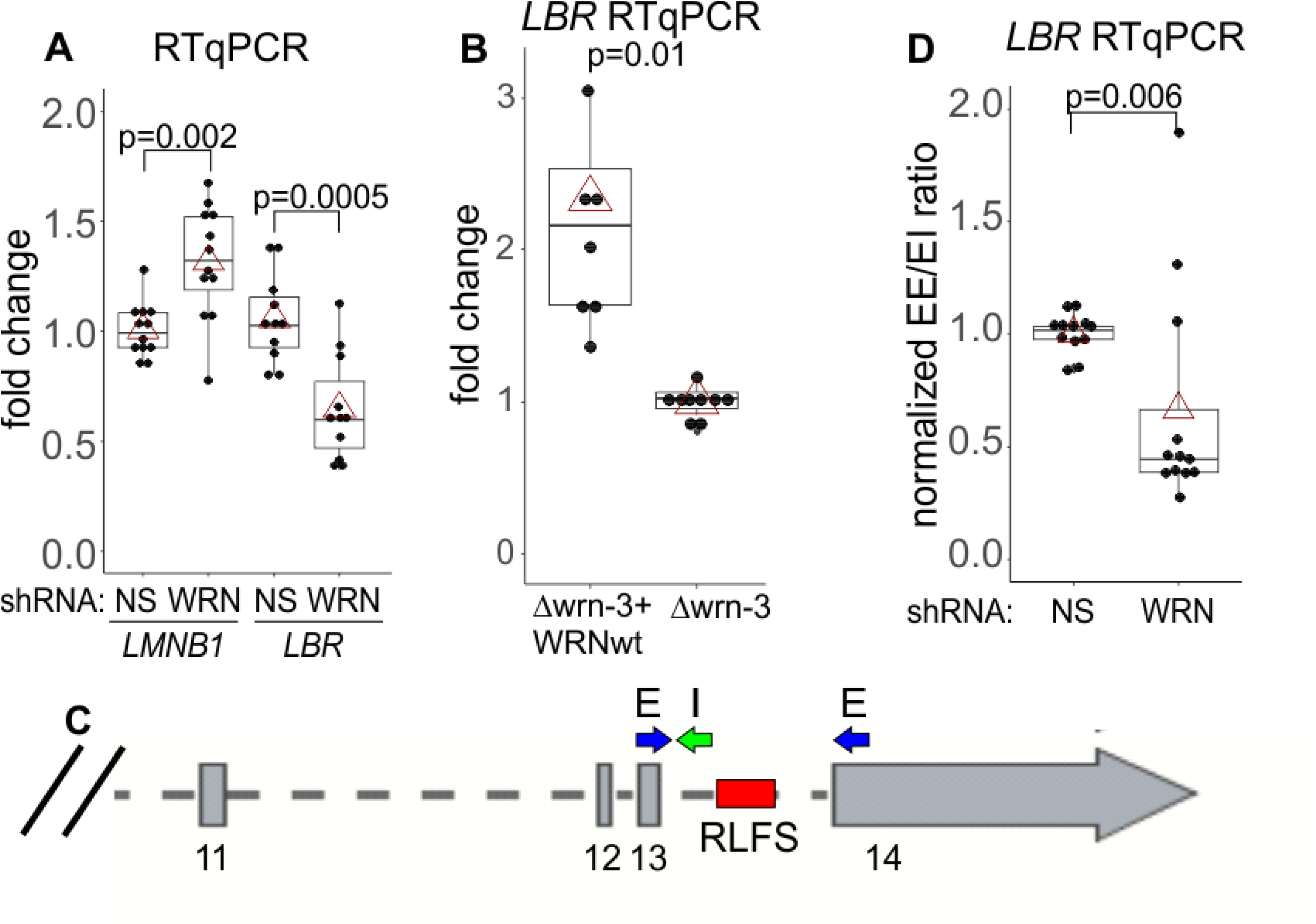
WRN facilitates transcription of the *LBR* gene. A, B) RT qPCR measurements of *LMNB1* and *LBR* gene transcripts in the WI38hTERT expressing the indicated shRNAs (A) or in Δwrn-3 cells with or without WRN complementation (B). C) A schematic of the 3’ end of the *LBR* gene with locations of the qPCR primers and an intronic R-loop forming site (RFLS). Exon numbers are below the gene. D) RT qPCR measurements of a ratio between the spliced LBR mRNA (EE, no intron 13) and the mRNA containing intron 13 (EI). The data are from WI38hTERT expressing the indicated shRNAs. Graphs in A, B, and D summarize three independent experiments each, with technical triplicates. P values were calculated in paired t-tests carried out on ΔΔ Cq values.

Notably, the *LBR* primer pair that we used maps to exons 13 and 14, the two last exons of the gene, as it is designed to detect a full-length, spliced mRNA (FIG.9C). The approx. 1Kb-long intron 13 between these exons contains an R-loop forming sequence (hg38_RLFS range=chr1:225403587-225404027), recognized as a high confidence RFLS by R-loopBase (Lin et al., 2021), R-loopDB (Jenjaroenpun et al., 2015)), and RLBase (Miller et al., 2022). Intron 13 RFLS is one of six such sequences in the introns of the *LBR* gene, all of which were experimentally verified to carry R loops *in vivo* as summarized in the above-mentioned databases. Moreover, the non-transcribed strand of the intron 13 RLFS is also rich in G4 motifs (chr1:225,403,870-225,404,030), which are recognized as G4-forming sites by G4Hunter Seeker (Bedrat et al., 2016). Since WRN can unwind R-loops and G-quadruplexes (Suzuki et al., 1997; Suzuki et al., 1999), we speculated that without WRN these secondary structures may be upregulated in *LBR* intron(s). Conceivably, this may downregulate splicing (Georgakopoulos-Soares et al., 2022), thus reducing the amount of the spliced mRNA detectable in our qPCR. We repeated RT-qPCRs measuring both the spliced and the intron 13-containing transcripts (FIG.9D). Indeed, the ratio between the levels of qPCR products detecting the spliced (EE in FIG.9) and the unspliced (EI in FIG.9) cDNAs was about two-fold lower in WRN-depleted cells, suggesting that at least the intron 13 of the transcript may be removed less efficiently. Translation of the intron 13-containing mRNA will result in a C-terminal 100 amino acid truncation of the LBR protein.

### Replication of SATII is affected by WRN depletion

We next asked whether or not WRN’s effect on heterochromatin maintenance is linked to the role of the protein in replication. Heterochromatin may impose unique demands on replication machinery. On the one hand, traversal of its compact structure may be a challenge (Chagin et al., 2019). On the other hand, some CH may be difficult to replicate due to the repetitive nature of its underlying DNA, which includes inverted repeats (Black and Giunta, 2018; Shastri et al., 2018; Zeller and Gasser, 2017). Indeed the SATIII class of satellites of the PCH was shown to replicate at a lower fork speed than the global genomic average (Mendez-Bermudez et al., 2018). WRN is known to support replication of problematic regions such as fragile sites (Drosopoulos et al., 2015) and inverted repeats (van Wietmarschen et al., 2020). Thus, it is possible that satellite replication may be differentially affected by WRN absence.

WRN role in replication of any of the satellite repeats has not been studied, and likewise the replication of SATII class of satellites, which comprise 1% of human genome (Altemose, 2022) is not well characterized. We looked at the replication of the SATII satellites using a combination of fiber FISH and maRTA, which allowed to see SATII arrays (green, FIG.10A) as well as replication tracks of CldU incorporation (red, FIG.10A) in stretched DNA fibers. SATII arrays are on average long enough to encompass a fork progressing for 30 min at an average genomic rate (FIG.10B), however, replication track lengths enclosed within SATII tracks (C_SAT_ tracks) were still significantly shorter than the global genomic average (C_GEN_ in FIG.10B). In WRN-deficient cells, C_GEN_ values were somewhat lower than in the control, highlighting a mildly reduced fork progression, as expected (Rodriguez-Lopez et al., 2002; Sidorova et al., 2013). Remarkably however, replication tracks within SATII arrays, C_SAT_, were somewhat longer than the C_SAT_ tracks of the control (FIG.10B). This translated into a reduced size of the difference between the genomic and the SATII-specific fork rates in WRN-depleted cells compared to the control (FIG.10D). These data suggest that, relative to each cell line’s average genomic fork progression, forks progress faster through the SATII heterochromatic arrays in WRN-deficient cells. This, together with our H3K9me3 ChIP-qPCR data (FIG.5), could imply that SATII heterochromatin is disrupted in WRN-depleted cells, affecting replication as a consequence.

**Figure 10.**
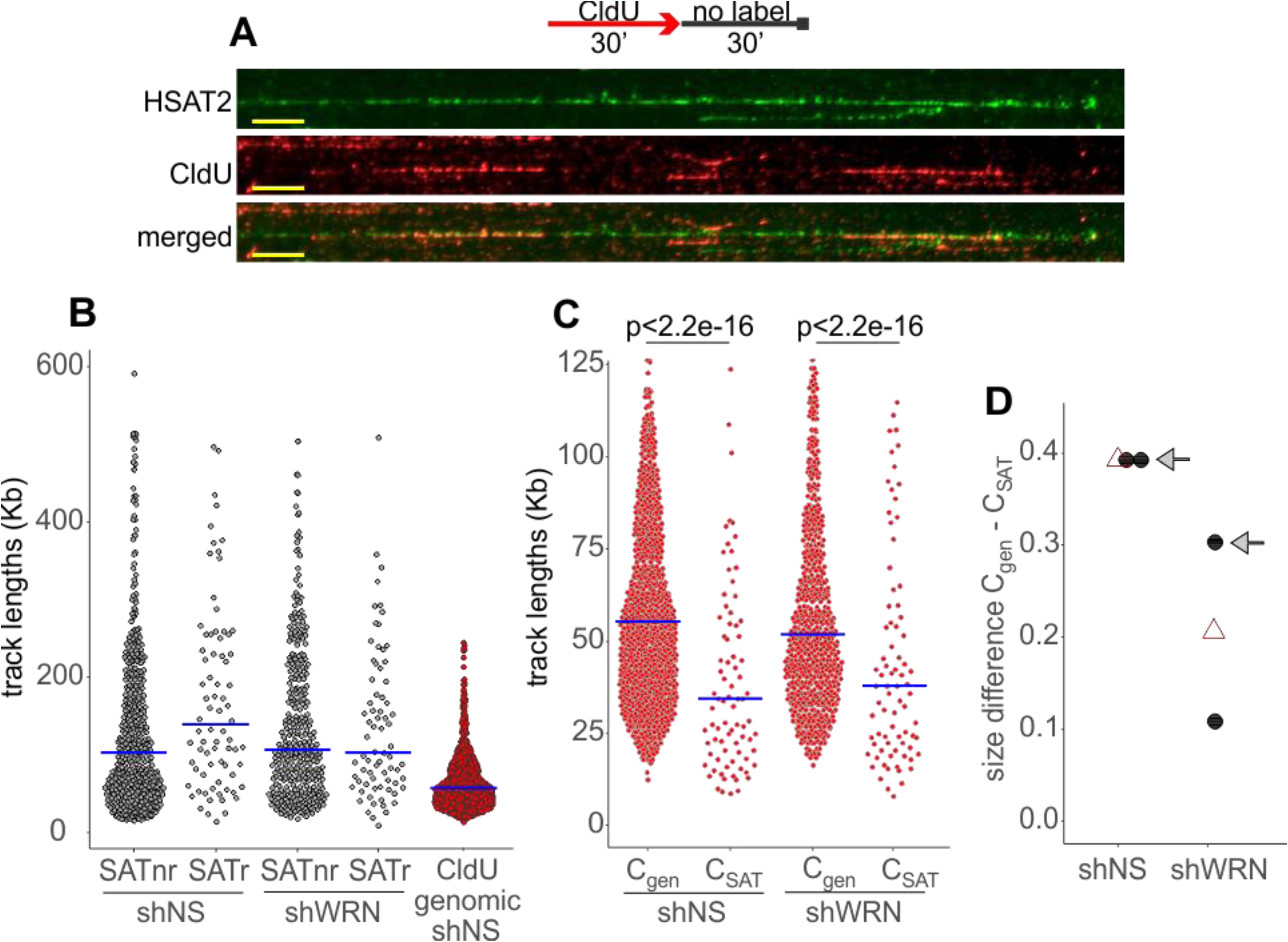
SATII satellite replication is affected by WRN deficiency in WI38hTERT. A) An example of a HSATII array (green) with CldU replication tracks (red) within it, as visualized by fiber FISH/maRTA. Scale, 10um. B) A labeling scheme and a plot of length distributions of HSATII tandems that do not (non-replicating, nr) or do have (replicating, r) CldU tracks within them. The length distribution of CldU tracks in total genomic DNA (excluding those within HSATII), is included in the same plot for comparison. B) Length distributions of nonreplicating (nr) and replicating (r) HSATII tracks. Genomic CldU tracks distribution is included for comparison. C) CldU tracks in the genome (C_gen_) and within HSATII arrays (C_SAT_). The plots are representative of two independent experiments. P values denote the probability with which a subset the size of n = n of C_SAT_ and a mean value equal to that of C_SAT_ can be randomly drawn from a C_gen_ dataset. Crossbars are distribution medians. C) Size of difference between the indicated CldU track length distributions in the control and WRN-depleted cells was determined by deriving Cliff’s delta statistic. Black dots are independent experiments and open triangles are means of the two experiments. The Cliff’s delta values derived from the experiment shown in (B) are marked by arrows. Smaller Cliff’s delta values indicate smaller difference between C_gen_ and C_SAT_.

To address this, we sought to quantify CH markers specifically on replicating DNA by using PLA with biotin-conjugated EdU, as previously (Lazarchuk et al., 2020; Lazarchuk et al., 2019). Cells were labeled with EdU and collected. The H3K9me3/EdU and LBR/EdU PLA signals were virtually undetectable under those conditions (not shown), however, we were able to detect a robust LMNB1/EdU PLA signal that was specific to EdU-positive cells (FIG.11A). As expected, EdU incorporation in WRN-depleted cells was somewhat lower (FIG.11B). LMNB1/EdU PLA signal was also consistently lower in WRN-depleted cells compared to the control (FIG.11C), but this differential could not be due merely to the lower EdU incorporation because it was maintained even after normalization by EdU intensity (FIG.11D). This result is consistent with the assumption that CH may be disrupted and thus may be less of an impediment to fork progression over SATII in WRN-depleted cells. In addition, it lends further support to the notion that WRN affects heterochromatin of proliferating cells.

**Figure 11.**
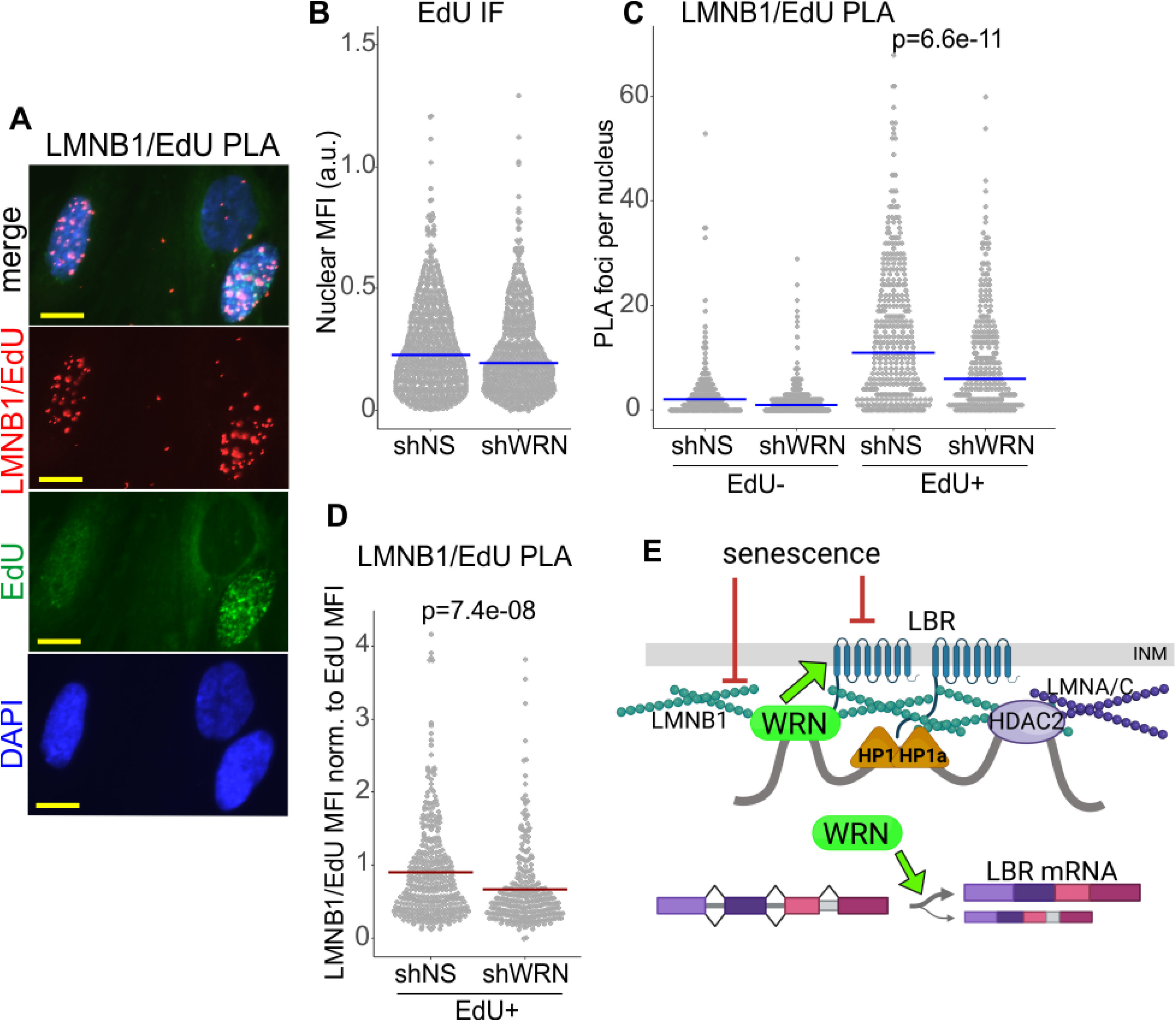
Lamin B1 association with replicating DNA is reduced in WRN-deficient cells. A) An example of a PLA analysis detecting Lamin B1 association with nascent DNA in situ, demonstrating specificity of the PLA signal to EdU+ positive cells. WI38hTERT cells (expressing non-targeting shRNA) pulse-labeled with 20uM EdU for 10 min and collected after a 10 min chase. EdU was clicked to a mixture of biotin-azide and Alexa488-azide at a molar ratio of 50:1. PLA was performed with antibodies against Lamin B1 and biotin. Scale bar, 10μm. B) Quantitation of EdU incorporation in WI38hTERT expressing the indicated shRNAs. C) Quantitation of LMNB1/EdU PLA in the same experiment as in (B). The datasets were subsetted into EdU- and EdU+ groups by applying a fixed cut-off EdU MFI value (e.g., 0.02 in (B)), below which cells were deemed EdU-. Lamin B1/EdU PLA PLA foci counts were then plotted separately for EdU- and EdU+ subsets. D) For each cell in EdU+ subpopulations, Lamin B1/EdU PLA foci counts were normalized to EdU MFI values and plotted. This normalization compensates for a somewhat lower level of EdU incorporation typically seen in WRN-depleted cells compared to the control (B). P values were calculated in Wicoxon tests. Blue crossbars are medians and red crossbars are means. The results of (B-D) represent two independent experiments, one of which included biological triplicates. E) A schematic of key findings.

## Discussion

Examination of CH in WRN-deficient proliferating, immortalized human fibroblasts suggests that several of its features are negatively affected in these cells, and the affected features are a subset of those also seen in senescing WRN-proficient cells. Most prominently, we see that WRN deficiency consistently reduces the associations involving Lamin B1 or LBR. In WI38hTERT but not GM639 background we also saw reduction of association of HP1α with KAP1. The apparent absence of this reduction in GM639 may be a detectability issue since not all cells in the complemented cell line express WRN. However, it is also possible that the effect of WRN on HP1α complexes is secondary to its effect on LBR and Lamin B1, and thus is more buffered against WRN manipulation. This is also consistent with the fact that both in WI38hTERT and GM639 WRN absence had no impact on association of HP1α with HDAC2. Further dissection of these dependencies will require identifying WRN mutants defective in specific associations.

Lamin B1 and LBR are responsible for tethering and compacting heterochromatin at the inner nuclear membrane, both via the HP1α protein and by directly interacting with other chromatin components (Douet et al., 2017; Hirano et al., 2012). In addition, Lamin B1 may participate in tethering of heterochromatin to the nucleolus and maintenance of nucleolar structure (Martin et al., 2009). Thus, the associations involving Lamin B1 or LBR in particular, together with the reductions of the repressive H3K9me3 mark on satellite repeats and of Lamin B1 association with nascent DNA, suggest that WRN deficiency reduces peripheral and possibly even nucleolar CH tethering in proliferating cells (FIG.11E).

Downregulation of LBR and Lamin B1 protein levels seen in the nuclei of WRN-deficient cells is likely sufficient to explain this reduction. Importantly, LBR and Lamin B1 downregulation is observed in senesced cells (Dillinger et al., 2017; En et al., 2020b; Kim, 2023; Lenain et al., 2015; Lukášová et al., 2018; Sadaie et al., 2013) and in some cell types is in fact sufficient to trigger senescence or SASP (En et al., 2020a; Herman et al., 2021; Lämmerhirt et al., 2022; Shimi et al., 2011). Furthermore, LBR downregulation was shown to also reduce Lamin B1 levels (Lukášová et al., 2017). Thus, by reducing the level of LBR, WRN deficiency may chronically compromise heterochromatin tethering and sensitize cells to senescence-promoting stimuli that target LBR and Lamin B1.

The above findings may suggest that WRN is an upstream regulator of LBR and Lamin B1. Though we were not able to see physical proximity of WRN to LBR or Lamin B1 with the antibodies used in this study, prior research (Zhang et al., 2015) and our findings of HP1α, HDAC2, WRN associations nevertheless indicate that WRN is also a physical constituent of the protein assemblies of heterochromatin. Furthermore, these WRN-involving assemblies are affected by senescence programs. We envision that WRN, HDAC2, and HP1α are proximal within the context of compacted heterochromatin, where their mutual proximity is dependent on its tethering to LBR and Lamin B1. Loss of this tethering in senescence results in separation of these proteins. Alternatively, senescence programs may target these interactions through posttranslational modifications of the participating proteins, including WRN (FIG.11E).

LBR reduction upon WRN loss has been noted before (Zhang et al., 2015) though no mechanism was explored. LBR abundance in the cell is controlled at the protein and RNA levels by multiple mechanisms that are deployed during differentiation (Solovei et al., 2013; Tiago et al., 2021; Wesley and Levy, 2023) or senescence (Arai et al., 2016; Herman et al., 2021). Our data suggest at least two ways by which WRN loss can compromise Lamin B1 and/or LBR-containing assemblies. First, we detected approximately a two-fold reduction of the LBR transcript in WRN-deficient compared to WRN-proficient cells. These data align with and confirm the LBR data in the transcriptomic datasets generated for isogenic WRN-depleted and control NHDFs (Tang et al., 2016) and in WRN KO vs. control MSCs (Zhang et al., 2015). Tang et al (Tang et al., 2016) observed that many of the genes whose transcription is affected by WRN contain potential G4-forming sequences at TSS and first introns, and proposed that WRN controls G4-quadruplex levels at these genes. Though the G4-rich RFLS in the intron 13 of *LBR* was not noted by these authors, *LBR* did make the list of potential G4-dependent targets of WRN owing to the G4 motifs present at its TSS and first intron. Based on these and our data, we hypothesize that in the absence of WRN an elevated R-loop and/or an RNA or DNA G-quadruplex may negatively affect splicing of the *LBR* transcript. This mechanism would be distinct from the recently uncovered suppression of the LBR transcript by the micro-RNA miR-340-5p that is induced in senescing WI38 cells (Herman et al., 2021). Indeed transcriptomic data did not show elevated miR-340-5p in WRN-deficient fibroblasts (Tang et al., 2016). Importantly, if translated, the unspliced intron 13 of LBR mRNA will terminate the LBR ORF prematurely at amino acid 521, and a similar truncation (at aa534) is known to cause Greenberg skeletal dysplasia and was shown to result in displacement of LBR into nucleoplasm and its proteasome-dependent degradation (Tsai et al., 2016). This, together with a possibility that the intron 13-containing mRNA may undergo nonsense-mediated decay, provides a plausible mechanism for reduction of total LBR in WRN-deficient cells.

WRN loss also reduces enrichment of the LBR protein in the nucleus, an effect that did not require WRN helicase or exonuclease activities, and has not been documented before. Subcellular distribution of LBR is dynamically regulated through cleavage, turnover and post-translational modification (Arai et al., 2016; Mimura et al., 2016). Phosphorylation of the arginine and serine-rich (RS) domain of LBR by the SRPK1 kinase leads to a more cytoplasmic distribution of the protein (Mimura et al., 2016). Oligomerization of LBR can be a factor in its retention at INM as well as in facilitating compaction of LBR-tethered heterochromatin, and hyperphosphorylation of the RS domain may disrupt oligomerization (Nikolakaki et al., 2017). In turn, subcellular redistribution of LBR can affect its stability, explaining the reduction of total LBR in WRN-depleted cells. These mechanisms can plausibly require WRN protein-protein interactions but not its enzymatic activity. Another possible mechanism can be derived from the earlier studies where WRN, via its interactor WHIP1, was found associated with the NUP proteins NUP107 through NUP160 of the nuclear pore complex, NPC (Li et al., 2013). Since NUP107 supports LBR localization to the INM (Mimura et al., 2016), it is possible that WRN absence can affect LBR localization via NPC activity or density.

We did not detect downregulation of the LMNB1 transcript in WRN-depleted cells, in contrast to the Tang et al study (Tang et al., 2016). This may be due to a difference in cell line backgrounds, assuming that more than one WRN-dependent mechanism may affect LMNB1 level. In our case, a previously mentioned dependence of the LMNB1 protein level on LBR (Lukášová et al., 2017) can explain our findings of LMNB1 reduction in WRN-depleted cells, though further experiments are needed to understand its mechanism.

One paradoxical result obtained in our study is that WRN does not seem to facilitate replication fork progression on SATII arrays. We and others previously demonstrated that WRN loss causes slower fork progression globally in the genome and in particular in the difficult to replicate loci such as fragile sites (Drosopoulos et al., 2015; Rodriguez-Lopez et al., 2002; Sidorova et al., 2013). Therefore, given that satellite repeats are viewed as potentially difficult to replicate and indeed display a significantly slower fork progression rate compared to the genomic average ((Mendez-Bermudez et al., 2018) and FIG.10), we anticipated that WRN loss would have a disproportionately large negative effect on fork progression on SATII arrays. This turned out not to be the case. It is possible that the protein complexes in charge of maintaining satellite repeat replication specifically exclude WRN. However, given the evidence of physical presence of WRN at SATII (Zhang et al., 2015), and the evidence of its presence in the two nuclear zones to which satellites localize, the nuclear periphery and the nucleolus (Li et al., 2013; Marciniak et al., 1998), this possibility seems less likely. Instead, we propose that CH is a factor that reduces the rate of fork progression on SATII. A negative impact of heterochromatin on replication fork rate is supported by the previous findings in which global loss (Lana et al., 2012) or global gain of heterochromatin (Igarashi et al., 2023; Kurashima et al., 2020) caused, respectively, fork acceleration or fork slowing/stalling. Our findings that in WRN-deficient cells replicating DNA has less Lamin B1 associated with it, and SATII repeats carry less H3K9me3 and are derepressed support the idea that CH is perturbed. If so, replication fork progression through it may indeed be expedited in WRN-deficient cells though it remains to be clarified how an increase in SATII transcription does not appear to interfere with replication.

In conclusion, our study highlights WRN as a contributor to the integrity of constitutive heterochromatin and points at the altered levels and distribution of LBR as a mediating mechanism. Our demonstration of this role of WRN in immortalized cells should facilitate its further mechanistic analyses as immortalized cells are a more utilizable model compared to limited-lifespan cells. The results also raise a question of whether the differential utilization of LBR as a heterochromatin tether by different cell types (Buxboim et al., 2017; Solovei et al., 2013; Tiago et al., 2021) can help to explain cell lineage-specific differentials in the severity of WRN deficiency, a phenomenon that is referenced in a definition of the Werner syndrome as a *segmental* progeria (Martin, 1985; Tsuge and Shimamoto, 2022). During iPS cells differentiation into mesoderm, LBR distribution changes from the predominantly nuclear to the mixed nuclear/cytoplasmic pattern (Wesley and Levy, 2023), which is reminiscent of the effect on LBR distribution produced by WRN depletion. Incidentally, it is the mesenchymal lineage that is most markedly affected by WRN loss (Tsuge and Shimamoto, 2022). It is tempting to speculate that lineages that are either more dependent on LBR and/or more rate-limiting for LBR may be more severely impaired by WRN loss. Further research is necessary to test this hypothesis and to fully understand the details of the functional interaction between WRN and LBR.

## Acknowledgements

We are grateful to Dr. Monnat for support and discussions and Dr. Weiliang Tang for a gift of the doxycycline-inducible WRN shRNA and control constructs. This work was supported by the NIH grants R01GM115482 to J.S. and R01CA210916 to J.O.

