## Supplementary figures and images for "Werner syndrome RECQ helicase participates in and directs maintenance of the protein complexes of constitutive heterochromatin in proliferating human cells"

### Supplemental Figures

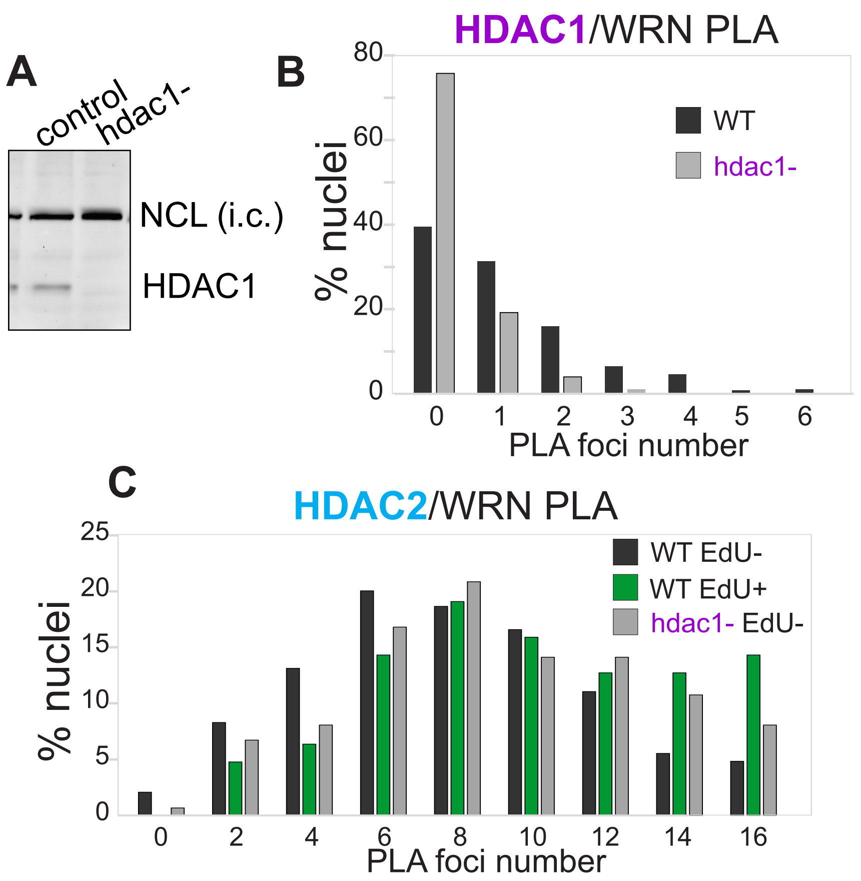

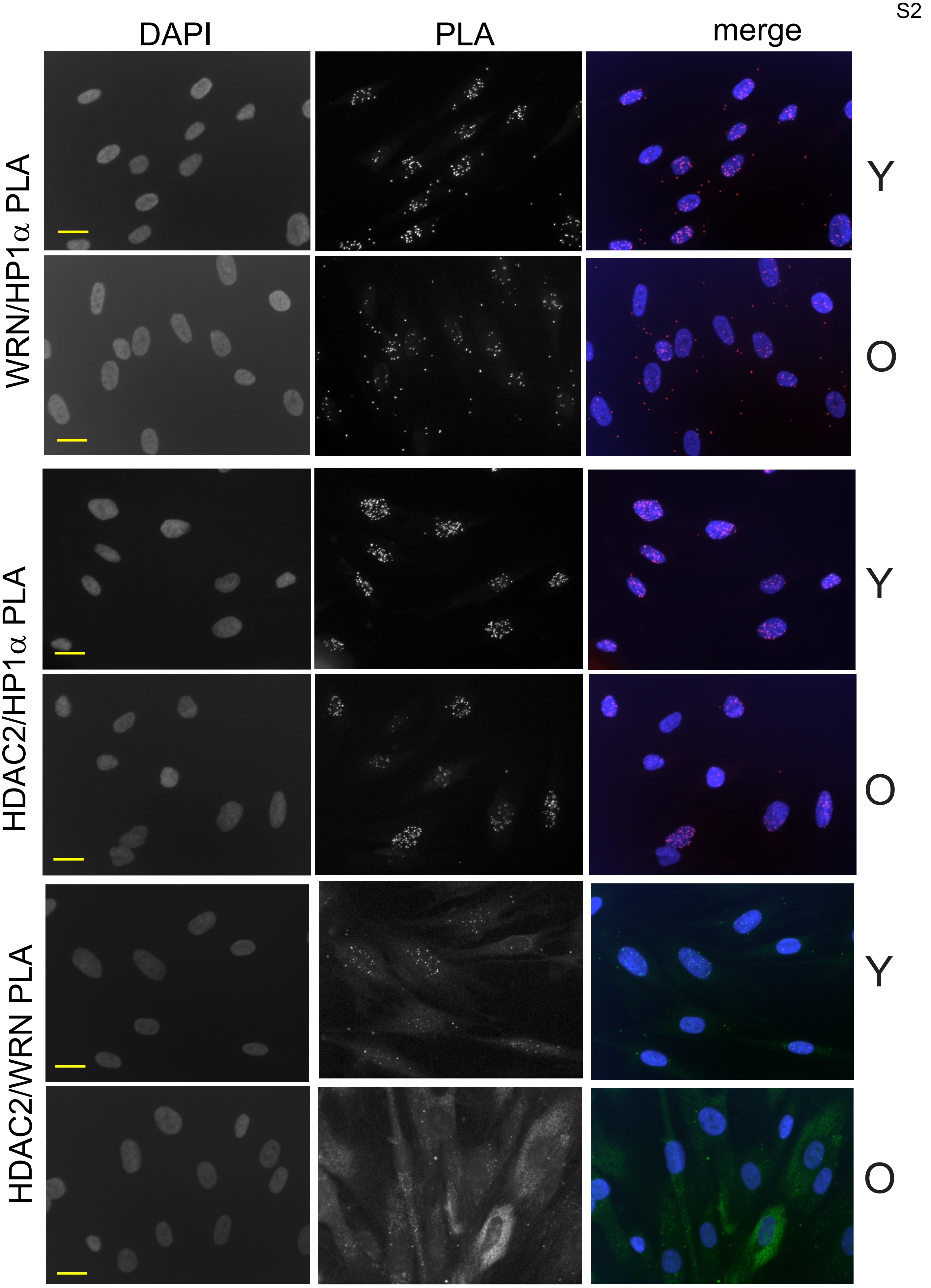

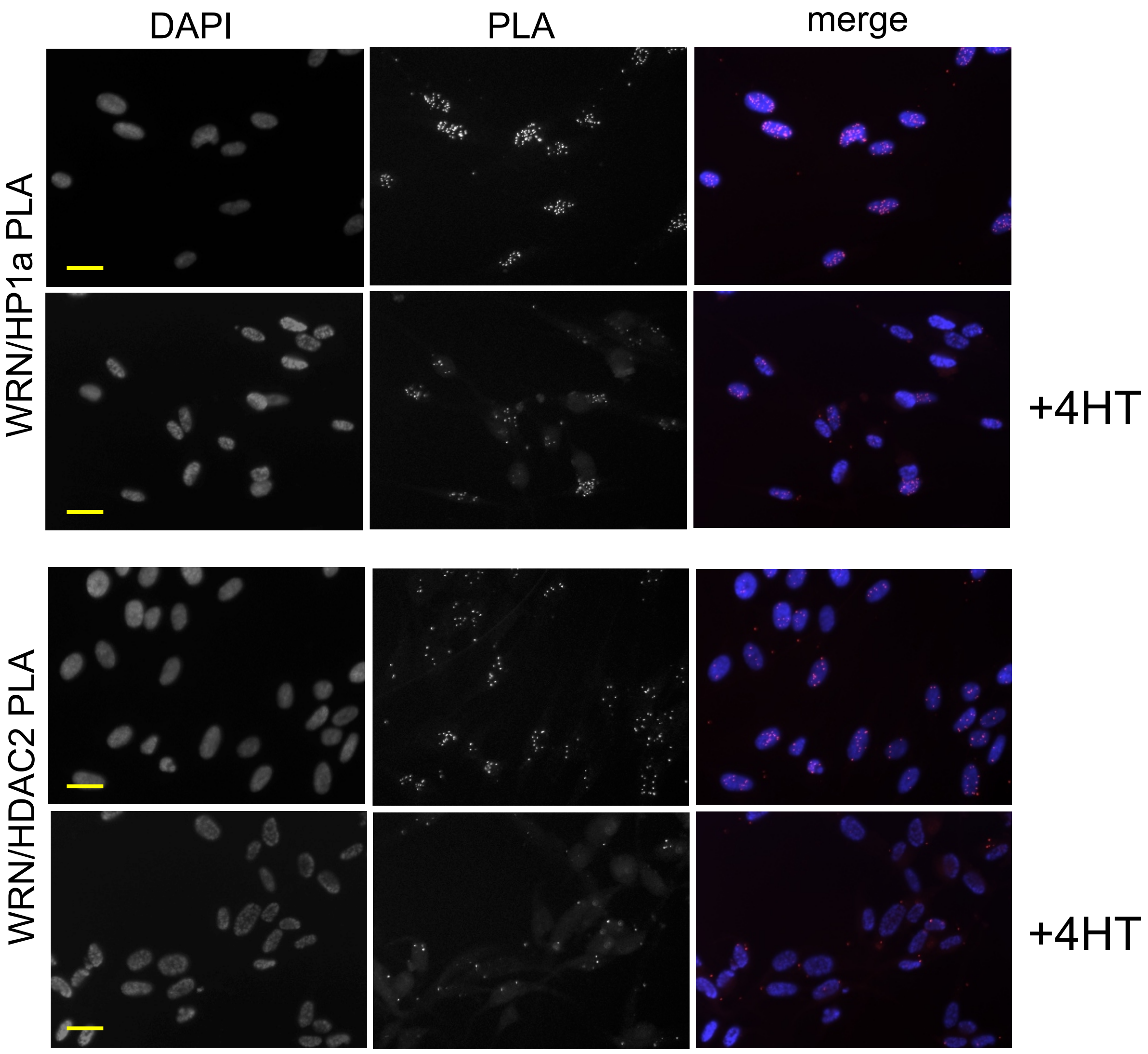

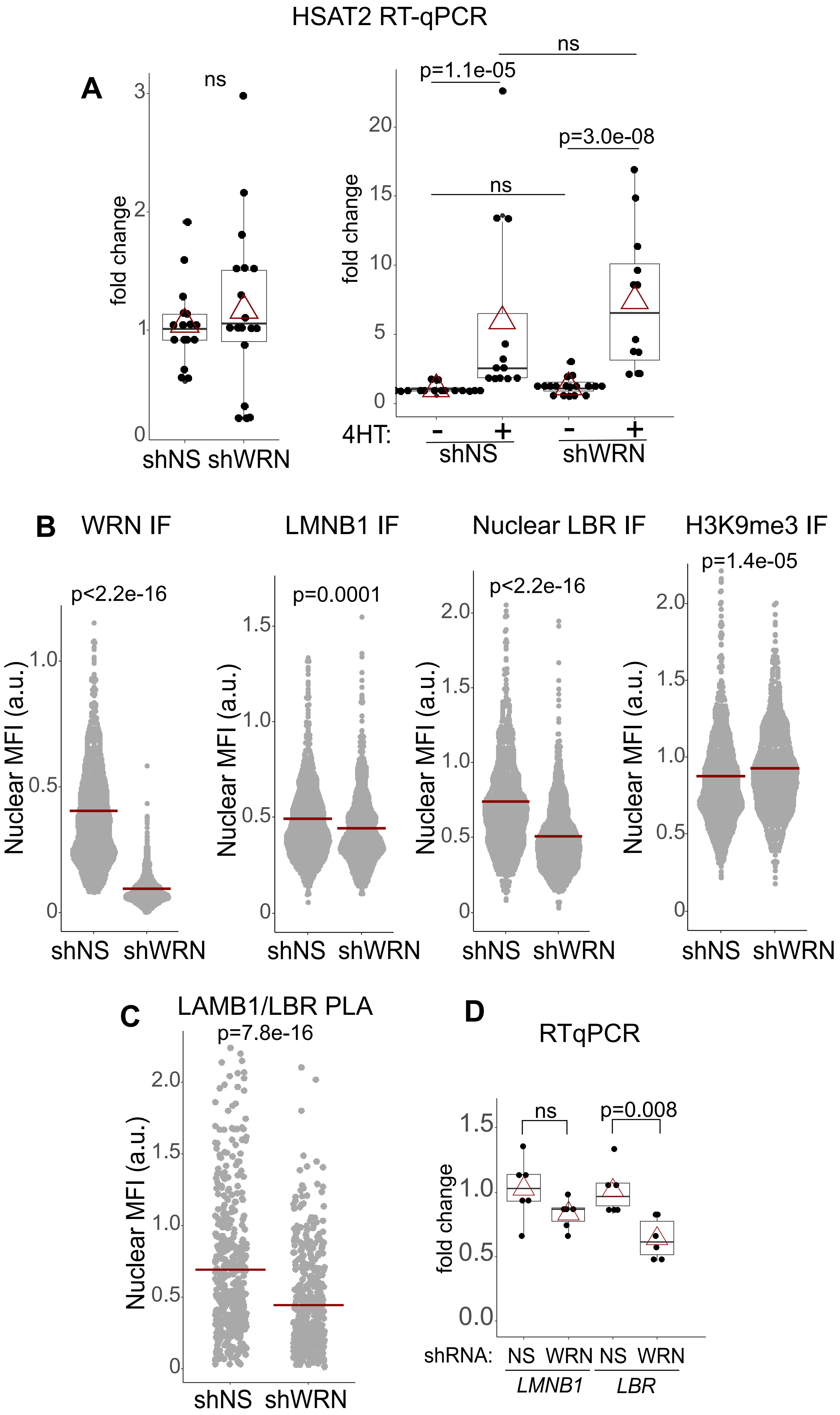

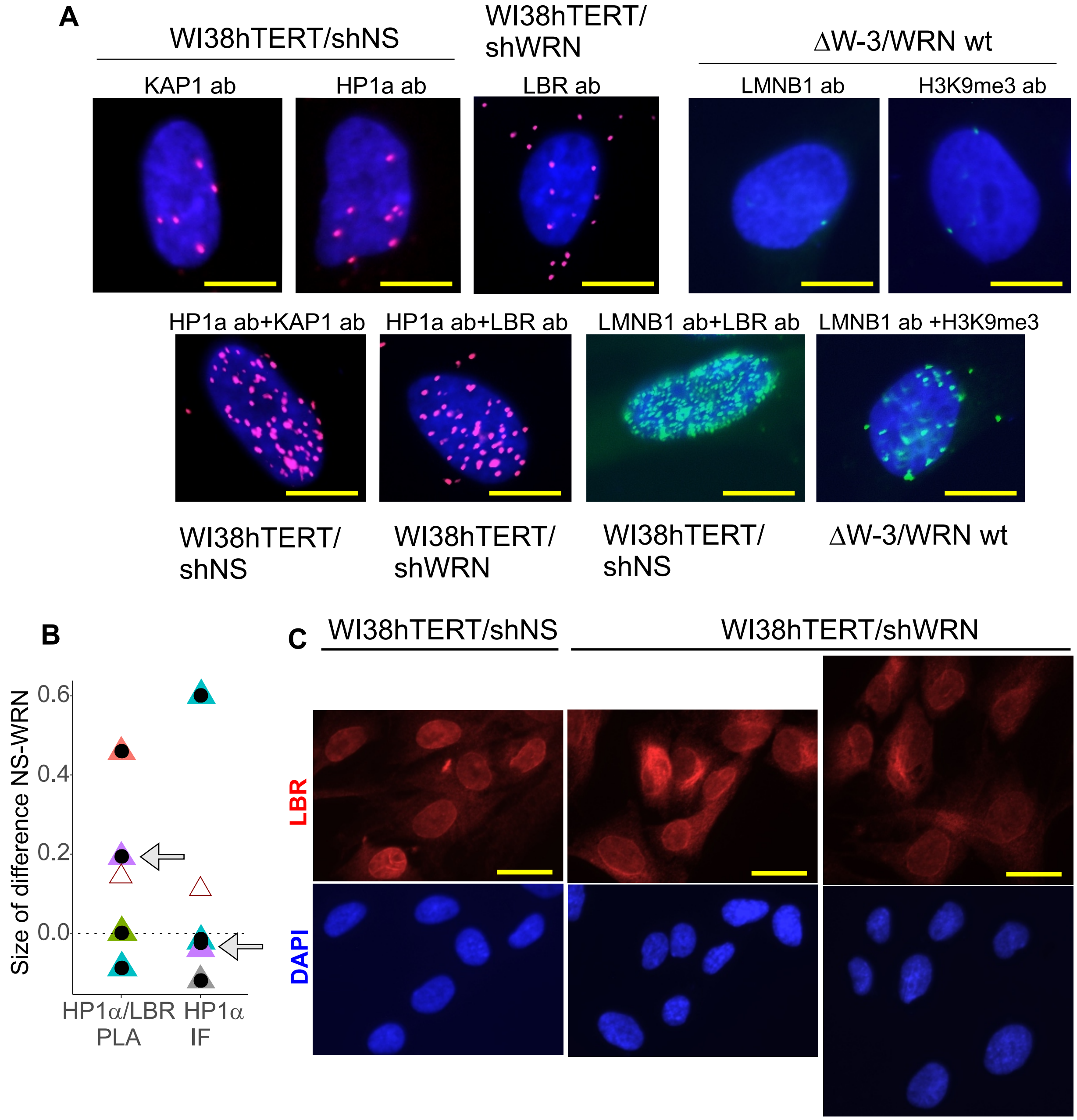

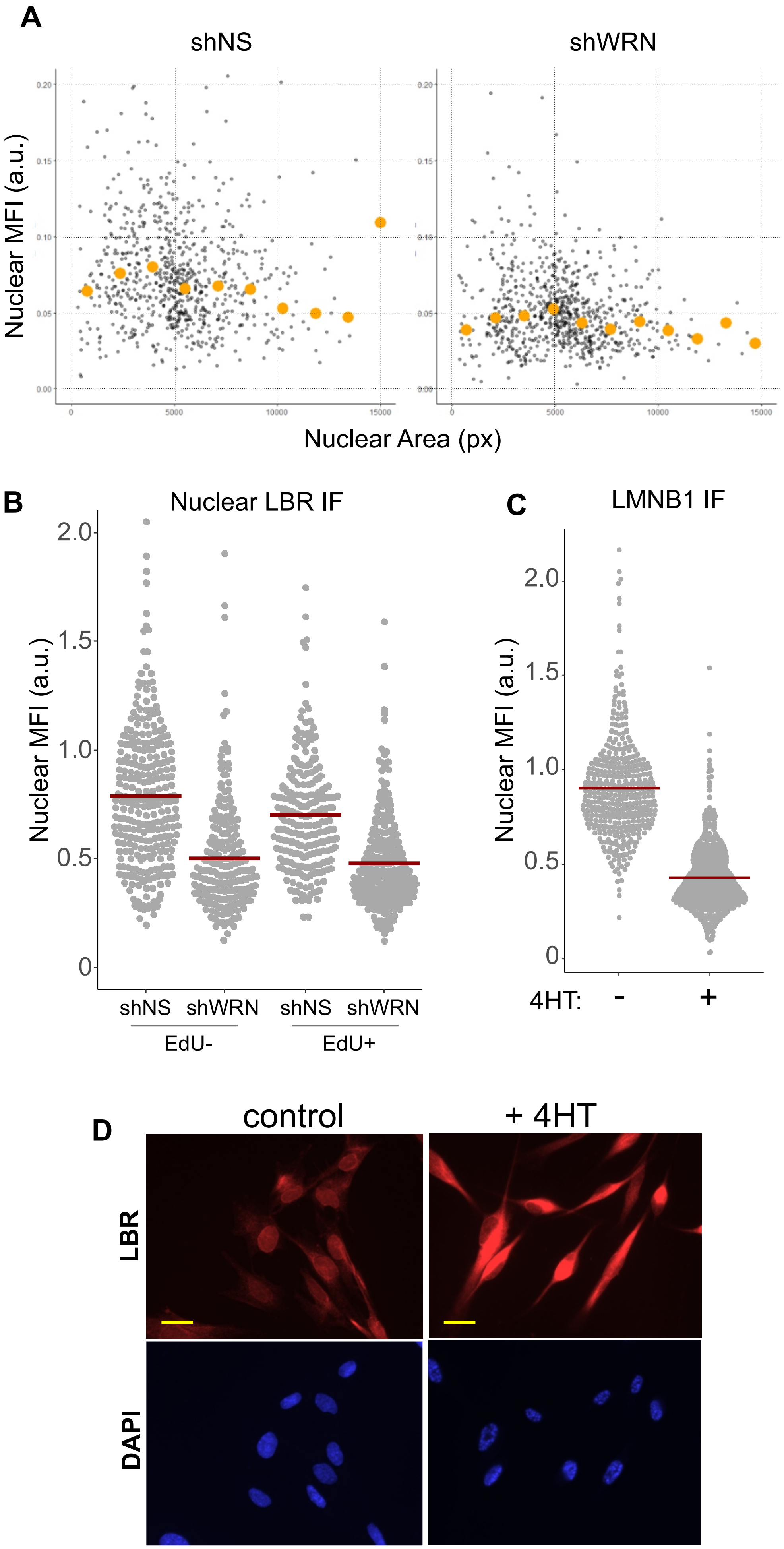
